# Computations and neural dynamics of audiovisual causal and perceptual inference in schizophrenia

**DOI:** 10.1101/2023.08.06.550662

**Authors:** Tim Rohe, Klaus Hesse, Ann-Christine Ehlis, Uta Noppeney

**Author notes:** **Corresponding author:** Tim Rohe **Email:**. **Author contributions:** TR and UN analyzed the data. TR and UN wrote the manuscript and KH as well as ACE critically reviewed it. TR and ACE conceived the experiment and TR as well as KH collected the data.

## Abstract

Hallucinations and perceptual abnormalities in psychosis are thought to arise from imbalanced integration of prior information and sensory inputs during perceptual inference. In this study, we combined psychophysics, Bayesian modelling and electroencephalography (EEG) to investigate potential changes in perceptual and causal inference in medicated individuals with schizophrenia when exposed to audiovisual sequences with varying numbers of flashes and beeps from either common or independent sources. Our findings reveal that individuals with schizophrenia, like their healthy controls, balance sensory integration and segregation in line with Bayesian causal inference rather than resorting to simpler heuristics. Both groups showed comparable weighting of prior information regarding the signals’ causal structure, with the schizophrenia group slightly overweighting prior information about the number of flashes or beeps. At the neural level, both groups computed Bayesian causal inference through dynamic encoding of perceptual estimates that segregate and flexibly combine audiovisual inputs. In conclusion, our results demonstrate that the computational and neural mechanisms of multisensory perceptual and causal inference remain remarkably intact in medicated individuals with schizophrenia during flash-beep scenarios.

## Introduction

Hallucinations - percepts in the absence of sources in the external world - are a hallmark of psychotic disorders such as schizophrenia. Individuals with psychosis may for instance hear voices or see people that are not present in their environment. The computational and neural mechanisms that give rise to these perceptual abnormalities remain unclear. Increasing research is guided by the notion that perception relies on probabilistic inference based on two distinct sources of information, observers’ top-down prior beliefs and bottom-up noisy sensory signals (Huys et al., 2015; Knill & Pouget, 2004; Ma, 2019; Noppeney, 2021; Parr et al., 2018; Pouget et al., 2013; Vilares et al., 2012). According to Bayesian probability theory, the brain should combine these two sources of information weighted according to their relative precisions (i.e. inverse of noise or variance), with a greater weight given to more reliable information. Hallucinations may thus arise from abnormal weighting of prior beliefs and incoming sensory evidence (Adams et al., 2013; Fletcher & Frith, 2009). Yet, while some studies have reported stronger or overly precise priors (Benrimoh et al., 2023; Cassidy et al., 2018; Powers et al., 2017; Teufel et al., 2015), others have associated psychosis with reduced or near-normal prior influences (Jardri et al., 2017; Schmack et al., 2017; Valton et al., 2019; Weilnhammer et al., 2020). These inconsistent findings raise the possibility that psychosis may either increase or decrease the precision or weight of different types of priors (Jardri et al., 2017; Schmack et al., 2017). While the weight of priors about simple features (e.g. motion) is thought to decrease in psychosis (Valton et al., 2019), the weight of priors about semantic or related information may increase (Fletcher & Teufel, 2022).

The weighting of various pieces of information becomes even more complex when the brain is confronted with multiple sensory signals that may come from same or different causes. In the face of this causal uncertainty, the brain needs to infer whether two sensory signals – say the sound of a whispering voice and the sight of articulatory movements – come from a common source (e.g., a single speaker) and should hence be integrated or else be processed independently (e.g., in case of different speakers). Normative models of hierarchical Bayesian causal inference (Kording et al., 2007; Shams & Beierholm, 2022; Shams & Beierholm, 2010) account for this causal inference problem by explicitly modelling the causal structures that could have generated the sensory signals. In the case of a common source, signals are fused weighted in proportion to their relative sensory precisions (Ernst & Banks, 2002); in the case of separate sources, they are processed independently. Critically, the brain needs to infer the underlying causal structure from spatiotemporal, numeric, semantic or other statistical correspondence cues (Parise & Ernst, 2016; Rohe et al., 2019; Rohe & Noppeney, 2015b; Wallace et al., 2004). A final percept is computed by combining the fusion (i.e. common source) and segregation (i.e. separate sources) estimates weighted by the posterior probabilities of each causal structure, a decision strategy referred to as model averaging (for other decisional functions, see (Wozny et al., 2010). Bayesian causal inference thereby enables a graceful transition from integration for (near-) congruent auditory and visual signals to segregation for incongruent signals. Recent research has shown that the brain accomplishes Bayesian causal inference by dynamically encoding the segregation, fusion and the final perceptual estimate that accounts for the brain’s causal uncertainty along cortical pathways (Aller & Noppeney, 2019; Cao et al., 2019; Rohe et al., 2019; Rohe & Noppeney, 2015a). Multisensory perceptual inference is thus governed by two sorts of priors, observers’ perceptual priors about environmental properties (e.g., number of signals) as well as a causal prior (or “binding tendency”) about whether signals come from common or independent sources. While the former influences observer’s perceptual estimates directly, the latter does so indirectly by modulating the strength of cross-sensory interactions.

The intricacies of multisensory perception may explain the inconsistent findings regarding multisensory abnormalities in psychosis (de Gelder et al., 2003; Haß et al., 2017; Noel et al., 2018; Vanes et al., 2016); for a review see Tseng et al., 2015). For instance, the rate at which participants with schizophrenia experience the McGurk- or sound-induced flash illusions has been shown to be lower (de Gelder et al., 2003; Vanes et al., 2016; White et al., 2014), equal (Balz et al., 2016) or even higher (Haß et al., 2017) compared to healthy controls. These inconsistencies may arise from the complex interplay of an individual’s auditory and visual precisions, perceptual and causal priors, and decisional strategies, which may all be altered in psychosis. Bayesian modelling and formal model comparison moves beyond previous descriptive approaches by allowing us to dissociate these distinct computational ingredients (Bennett et al., 2019; Huys et al., 2016; Jones & Noppeney, 2021).

This psychophysics-EEG study investigated whether schizophrenia alters the computational and/or neural mechanisms of multisensory perception in a sound-induced flash illusion paradigm. In an inter-sensory selective attention task, patients with schizophrenia and age-matched healthy controls were presented with sequences of a varying number of flashes and beeps. We first assessed whether schizophrenia altered the computations of how observers combined auditory and visual signals into number estimates by comparing normative and approximate Bayesian causal inference (BCI) models. Next, we combined BCI modelling with multivariate EEG analyses to unravel the underlying neural mechanisms. Our results suggest that multisensory inference processes are remarkably preserved in our medicated SCZ patient cohort compared to healthy participants at the behavioral, computational and neural level.

## Results

### Experimental design and analysis

23 healthy controls (HC) and 17 observers with schizophrenia (SCZ) were included in the EEG analysis. HC were matched to SCZ individuals in sex, age and education (Supporting Tab. S1). Neuropsychological tests revealed comparable performances in attentional and executive functions across the two groups. Yet, HC had better memory recall and premorbid crystallized intelligence.

In a sound-induced flash-illusion paradigm, we presented HC and SCZ with flash-beep sequences and their unisensory counterparts. Across trials, the number of beeps and flashes varied independently according to a four (1 to 4 flashes) × four (1 to 4 beeps) factorial design (Fig. 1A, B). Thereby, the paradigm yielded numerically congruent or incongruent flash-beep sequences on four levels of audiovisual numeric disparity. In an inter-sensory selective attention task, observers reported either the number of beeps or flashes.

**Figure 1.**
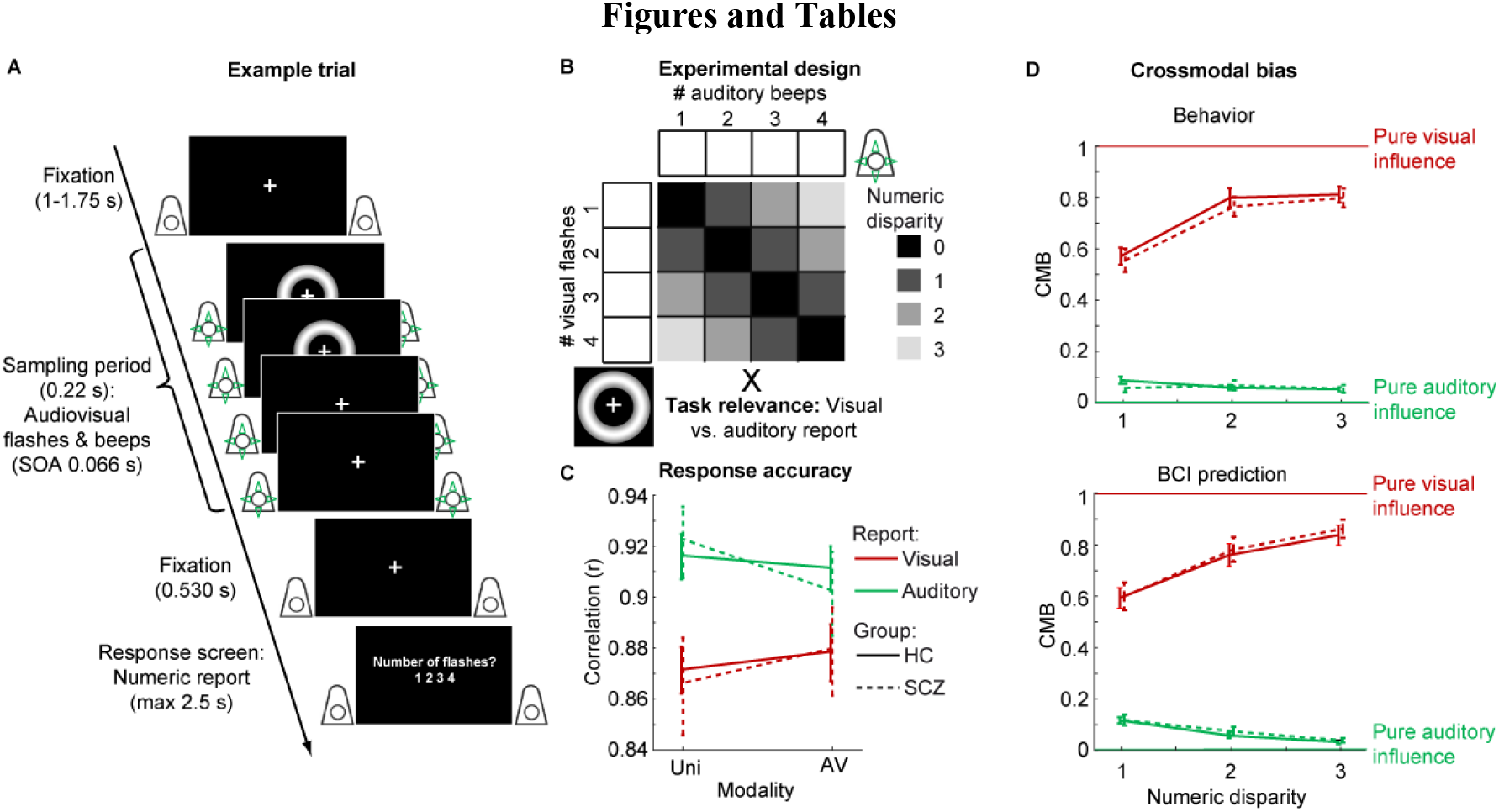
Example trial, experimental design and behavioral data. **(A)** Example trial of the flash-beep paradigm (e.g. two flashes and four beeps are shown) in which participants either report the number of flashes or beeps. **(B)** The experimental design factorially manipulated the number of beeps (i.e. one to four), number of flashes (i.e. one to four) and the task relevance of the sensory modality (report number of visual flashes vs. auditory beeps). We reorganized these conditions into a two (task relevance: auditory vs. visual report) × two (numeric disparity: high vs. low) factorial design for the GLM analyses of the audiovisual crossmodal bias. **(C)** Response accuracy (across-participants mean ± SEM; *n* = 40) is shown as a function of modality (audiovisual congruent conditions vs. unisensory visual and auditory conditions), task relevance (auditory vs. visual report) and group (HC vs. SCZ). **(D)** The audiovisual crossmodal bias (CMB; across-participants mean ± SEM; *n* = 40) is shown as a function of numeric disparity (1, 2 or 3), task relevance (auditory vs. visual report) and group (HC vs. SCZ). CMB was computed from participants’ behavior (upper panel) and from the prediction of the individually fitted BCI model (lower panel; i.e., model averaging with increasing sensory variances). CMB = 1 for purely visual and CMB = 0 for purely auditory influence.

### Behavior – scalar variability, response accuracy and crossmodal bias

Figure 2A shows the reported flash and beep counts in SCZ and HC as a function of the true flash and beep numbers. Both SCZ and HC progressively underestimated the increasing numbers of flashes and beeps (Supporting Fig. S1 and Tab. S2). This logarithmic compression and the increasing variances for greater number of flashes/beeps was similarly observed in both groups. It is in line with the known scalar variability of numerosity estimates (Dehaene, 2007; Gallistel & Gelman, 2000). Figure 2 also indicates that observers’ reported flash (resp. beep) counts were biased towards the concurrent incongruent beep (resp. flash) number in the ignored sensory modality. We formally compared SCZ and HC in their ability to selectively estimate either the number of flashes or beeps (see Fig. 1C, Tab. 1) using a 2×2×2 mixed-model ANOVA, with factors group (SCZ vs. HC), stimulus modality (unisensory visual/auditory vs. audiovisual congruent), and task relevance (auditory vs. visual report) on response accuracies (n.b. no incongruent conditions were included in this ANOVA). No significant group differences or interactions with group were observed. Instead, Bayes factors provided substantial evidence for comparable accuracies of the flash and beep counts in HC and SCZ (i.e., BF_incl_ < 1/3, Tab. 1; see Supporting Fig. S2 for comparable response time results).

**Figure 2.**
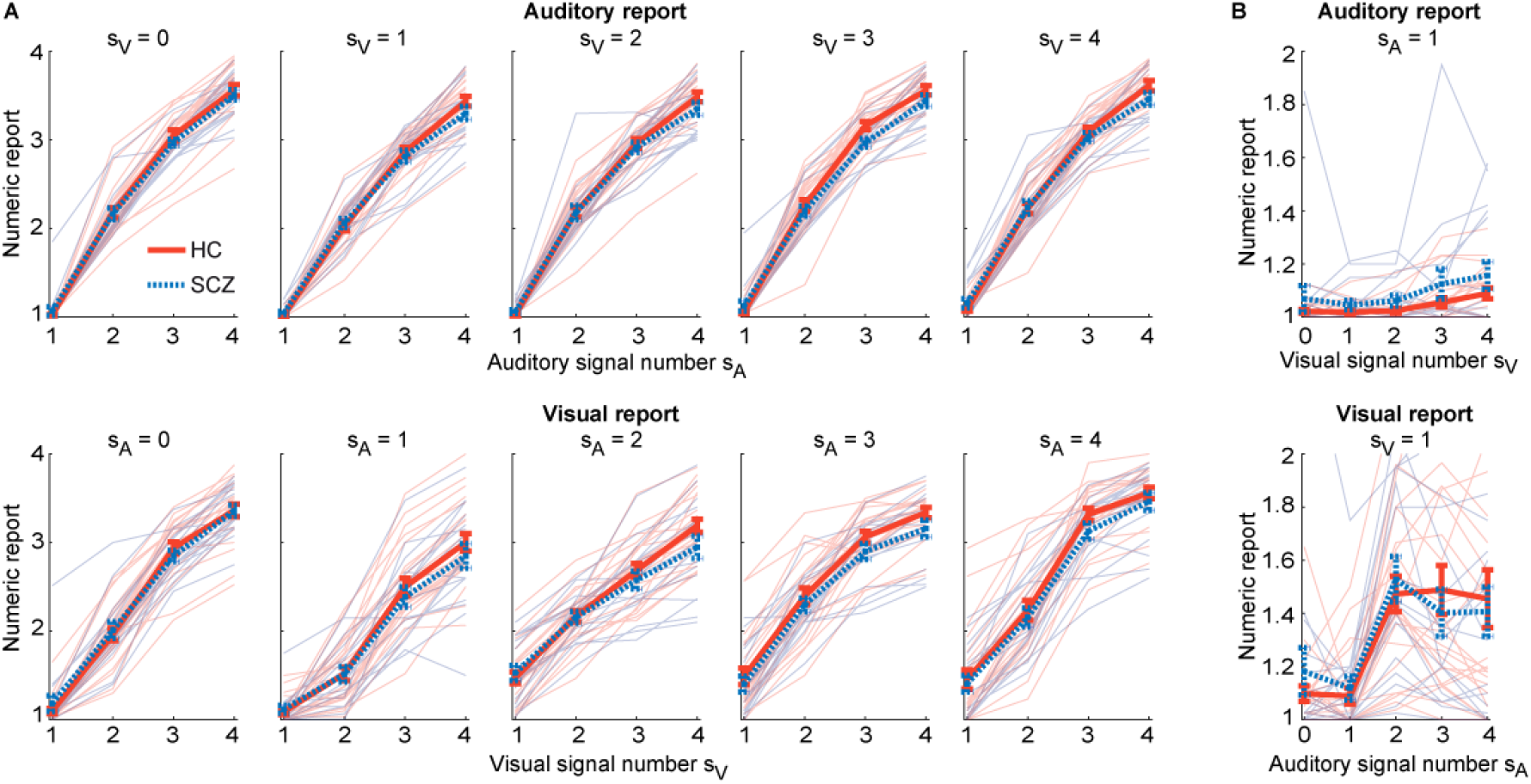
Distributions of numeric reports (across-participants mean ± SEM, n = 40) for HC and SCZ patients. **(A)** Upper panel: The auditory numeric reports plotted as a function of auditory signal number s_A_, separately for different visual signal numbers s_V_. Lower panel: The visual numeric reports plotted as a function of visual signal number s_V_, separately for different auditory signal numbers s_A_. **(B)** Auditory reports for a single beep as a function of visual signal number (upper panel) and visual reports for a single flash as a function of auditory signal number (lower panel).

**Table 1.** Main and interaction effects of task relevance (TR), modality and group on response accuracy and effects of task relevance, disparity and group on the crossmodal bias from classical and Bayesian mixed-model ANOVAs.

| | F | df1, df2 | p | part. $\eta^2$ | BF<sub>incl</sub> |
| --- | --- | --- | --- | --- | --- |
| **Response accuracy** |  |  |  |  |  |
| TR | 49.548 | 1,38 | <0.001 | 0.566 | >100 |
| Mod | 0.590 | 1,38 | 0.447 | 0.015 | 0.602 |
| Group | 0.006 | 1,38 | 0.939 | <0.001 | 0.233 |
| TR×Mod | 6.693 | 1,38 | 0.014 | 0.150 | 2.452 |
| TR×Group | 0.004 | 1,38 | 0.950 | <0.001 | 0.352 |
| Mod×Group | 0.625 | 1,38 | 0.434 | 0.016 | 0.186 |
| TR×Mod×Group | 1.691 | 1,38 | 0.201 | 0.043 | 0.145 |
| **Crossmodal bias** |  |  |  |  |  |
| TR | 633.673 | 1,38 | <0.001 | 0.943 | >100 |
| Disp | 92.312 | 1.7,65.7 | <0.001 | 0.708 | >100 |
| Group | 0.312 | 1,38 | 0.579 | 0.008 | 0.189 |
| TR×Disp | 90.781 | 1.6,61.4 | <0.001 | 0.705 | >100 |
| TR×Group | 0.075 | 1,38 | 0.786 | 0.002 | 0.068 |
| Disp×Group | 0.456 | 1.7,65.7 | 0.608 | 0.012 | 0.146 |
| TR×Disp×Group | 0.914 | 1.6,61.4 | 0.388 | 0.023 | 0.049 |

Note: TR = task relevance: auditory vs. visual report; Mod = modality: audiovisual congruent vs. unisensory; Group: HC vs. SCZ; Disp = absolute numeric disparity (1 vs. 2 vs. 3). Effects of the classical mixed-model ANOVA are Greenhouse-Geisser corrected if sphericity is violated.

As expected (Andersen et al., 2004), we observed a significant main effect of task-relevance and a task-relevance x modality interaction on response accuracies. Overall, observers estimated the number of beeps more accurately than the number of flashes. They also estimated the number of beeps better when presented alone than together with flashes, while the reverse was true for observers’ flash counts. Consistent with extensive previous research showing a greater temporal precision for the auditory than the visual sense (Burr et al., 2009), our results confirm that observers obtained more precise estimates for the number of beeps than for the number of flashes.

Based on Bayesian probability theory, observers should therefore assign a stronger weight to the more precise auditory signal when integrating audiovisual signals into number estimates, resulting in the well-known sound-induced flash illusion (Buergers & Noppeney, 2022; Rohe et al., 2019; Shams et al., 2000). Consistent with this conjecture, both SCZ and HC were more likely to perceive two flashes when a single flash was presented together with two sounds (i.e., fission illusion) and a single flash when two flashes were presented together with one sound (i.e., fusion illusion; see Supporting Fig. S3). We quantified these audiovisual interactions using the crossmodal bias (CMB) that ranges from pure visual (CMB = 1) to pure auditory (CMB = 0) influence (Figure 1D; n.b. the crossmodal can be computed only for numerically disparate flash-beep sequences). Consistent with the principles of precision-weighted integration (Ernst & Banks, 2002), the HC and SCZ’s flash reports were biased towards the number of auditory beeps (i.e., CMB < 1, t_39_ = −11.864, p < 0.001, Cohen’s d = - 1.876, BF_10_ > 100), again with no significant differences between the groups (t_38_ = 0.434, p = 0.667, d = 0.139, BF_10_ = 0.336). By contrast, we observed only a small but significant crossmodal bias for auditory reports towards the number of flashes in HC and SCZ (Fig. 1D; CMB > 0, t_39_ = 8.550, p < 0.001, d = 1.352, BF_10_ > 100) - again with no evidence for group differences (t_38_ = 0.448, p = 0.657, d = 0.143, BF_10_ = 0.337). Thus, both HC and SCZ assigned a greater weight to the temporally more reliable auditory sense.

An additional 2 × 3 × 2 mixed-model ANOVA with factors group (SCZ vs HC), audiovisual numeric disparity between beeps and flashes (1, 2 vs. 3) and task-relevance (auditory vs visual report) revealed that these crossmodal biases were significantly decreased at large numeric disparities, when signals most likely originate from different sources and should hence be segregated (i.e. task relevance × numeric disparity interaction, Tab. 1). At large numeric disparities, HC and SCZ were thus able to selectively report the number of flashes (or beeps) with minimal interference from task-irrelevant beeps (or flashes). This task relevance × numeric disparity interaction is qualitatively the key response profile predicted by Bayesian causal inference. Again, Bayes factors indicated substantial to strong evidence for comparable performance in SCZ and HC (i.e., BF_incl_ < 1/3 or even < 1/10, Tab. 1).

Overall, these GLM-based analyses of behavioral data (i.e. accuracy, crossmodal bias) suggest that both SCZ and HC combine audiovisual signals into number estimates qualitatively consistent with the principles of Bayesian causal inference. Both groups gave a stronger weight to the more reliable auditory signals leading to the well-known sound-induced flash illusions.

### Behavior – Bayesian modelling

To quantitatively assess whether HC and SCZ combine audiovisual signals according to shared computational principles, we compared 10 Bayesian models in a 5 × 2 factorial model space spanned by the modelling factors of ‘decision strategy’ (5 levels) and ‘sensory variance’ (2 levels) (Acerbi et al., 2018; Rohe & Noppeney, 2015b; Wozny et al., 2010). All models conformed to the basic architecture of the BCI model (Fig. 3).

**Figure 3.**
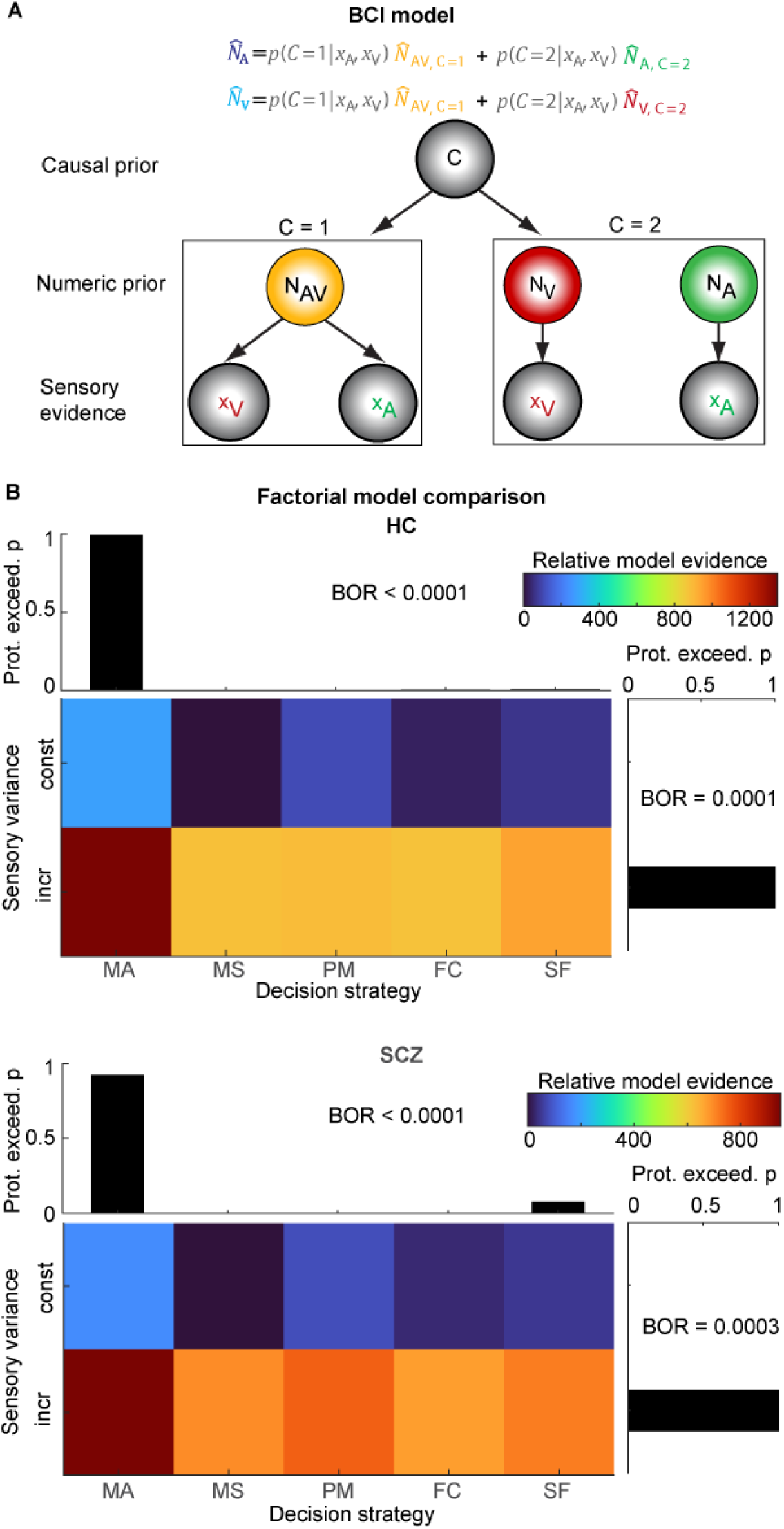
The BCI model and factorial model comparisons in HC and SCZ. **(A)** The BCI model assumes that audiovisual stimuli are generated depending on a causal prior (p_Common_): In case of a common cause (C=1), the “true” number of audiovisual stimuli (N_AV_) is drawn from a common numeric prior distribution (with mean μ_P_) leading to noisy auditory (x_A_) and visual (x_V_) inputs. In case of independent causes (C=2), the “true” auditory (N_A_) and visual (N_V_) numbers of stimuli are drawn independently from the numeric prior distribution. To estimate the number of auditory and visual stimuli given the causal uncertainty, the BCI model estimates the auditory or visual stimulus number (*N̂*_A_ or *N̂*_V_, depending on the sensory modality that needs to be reported). In the model-averaging decision strategy, the BCI model combines the forced-fusion estimate of the auditory and visual stimuli (*N̂*_AV,C=1_) with the task-relevant unisensory visual (*N̂*_V,C=2_) or auditory estimates (*N̂*_A,C=2_), each weighted by the posterior probability of a common (C = 1) or independent (C = 2) causes, respectively (i.e. *p*(*C* = 1| *x*_A_, *x*_V_) or p(C = 2| *x*_A_, *x*_V_)). **(B)** The factorial Bayesian model comparison (n = 40) of models with different decision strategies (model averaging, MA; model selection, MS; probability matching, PM; fixed criterion, FC; stochastic fusion, SF) with constant or increasing sensory auditory and visual variances, separately for HC and SCZ. The grayscale images show the relative model evidence (i.e., the Bayesian information criterion relative to the winning MA model with increasing sensory variances) for each model. The bar plots show the protected exceedance probability (i.e. the probability that a given model is more likely than any other model, beyond differences due to chance) for each model factor. The Bayesian omnibus risk (BOR) estimates the probability that factor frequencies purely arose from chance.

Along the factor ‘decision strategy’, we manipulated how observers combined the estimates formed under fusion (i.e. common source) and segregation (i.e. separate sources) assumptions into a final perceptual estimate. While growing research has shown that HC combine audiovisual signals according to the decision strategy of model averaging (Rohe et al., 2019; Rohe & Noppeney, 2015a, 2015b), we hypothesized that SCZ may resort to suboptimal strategies or even simpler heuristics such as applying a fixed threshold on audiovisual numeric disparity (Acerbi et al., 2018). In other words, rather than arbitrating between sensory integration and segregation according to the posterior probability of a common cause in a Bayesian fashion, SCZ may simply do so based on audiovisual spatial disparity.

In total, we compared the following five decisional strategies: i. Model averaging combined the segregation and fusion estimates weighted by the posterior probabilities of each causal structure. ii. Model selection selected the perceptual estimate from the causal structure with the highest posterior probability. iii. Probability matching selected either the fusion or segregation estimates in proportion to their posterior probability. iv. The fixed-criterion threshold model incorporates the simple heuristic of selecting the segregation estimate when the audiovisual numeric disparity exceeded a fixed threshold. v. Finally, the probabilistic fusion model employs the suboptimal strategy of selecting either the fusion or segregation estimates with a fixed probability that is estimated from observers’ responses (see methods for details and Acerbi et al., 2018). Additionally, we assessed along the modelling factor ‘sensory variance’ whether auditory and visual noise variances were constant or changed with the number of beeps/flashes as predicted by the scalar-variability of numerosity estimation (Dehaene, 2007; Gallistel & Gelman, 2000) and initial inspection of our data (see Fig. 2, Supporting Fig. S1).

Comparing all 10 models in this 2 × 5 factorial model space (Fig. 3 and Supporting Tab. S3) revealed that the model-averaging model with increasing sensory variances outperformed all other 9 models in both HC and SCZ observers. Furthermore, a between-group Bayesian model-comparison provided strong evidence that HC and SCZ individuals relied similarly on the five decision strategies, and in particular on model-averaging as the “winning” strategy, equally often (BF_10_ = 0.022). These Bayesian model-comparison results further support the notion that SCZ, like HC, performed multisensory perceptual and causal inference according to the same computational and decision strategies.

Having established that SCZ and HC combine audiovisual signals in line with Bayesian causal inference and read out the final estimate according to model averaging, we examined whether SCZ may overweight their prior about the signals’ causal structure (i.e. p_common_). Contrary to this conjecture, two-sample randomization tests on p_common_ did not reveal any significant difference between the two groups (Fig. 4 and Tab. 2). In the next step, we asked whether SCZ may rely more strongly on their priors about the flash/beep number relative to the sensory inputs as may be hypothesized based on a growing number of studies in unisensory perception (Benrimoh et al., 2023; Cassidy et al., 2018; Powers et al., 2017; Teufel et al., 2015). An over-reliance on prior information would be reflected in a greater precision (i.e. smaller variance; σ_P_) of numeric priors (i.e. *µ*_P_) in SCZ relative to HC. As shown in Figure 4 and Table 2, two-sample randomization tests indeed revealed a significantly smaller variance for the numeric prior in SCZ compared to HC, although the effect was small. This more precise numeric prior accounts for the fact that the numeric reports are centered more around the numeric prior mean in SCZ compared to HC (Tab. 2; cf. Fig. 2). Further, we observed a positive correlation between SCZ’s positive symptoms with their visual variance (r = 0.625, t_16_ = 3.101, p = 0.003, BF_10_ = 6.518; randomization test of the correlation). Consistent with previous research (Benrimoh et al., 2023; Cassidy et al., 2018; Powers et al., 2017; Teufel et al., 2015), this pattern of results is consistent with the general notion that SZC rely more on prior knowledge than new incoming visual evidence.

**Figure 4.**
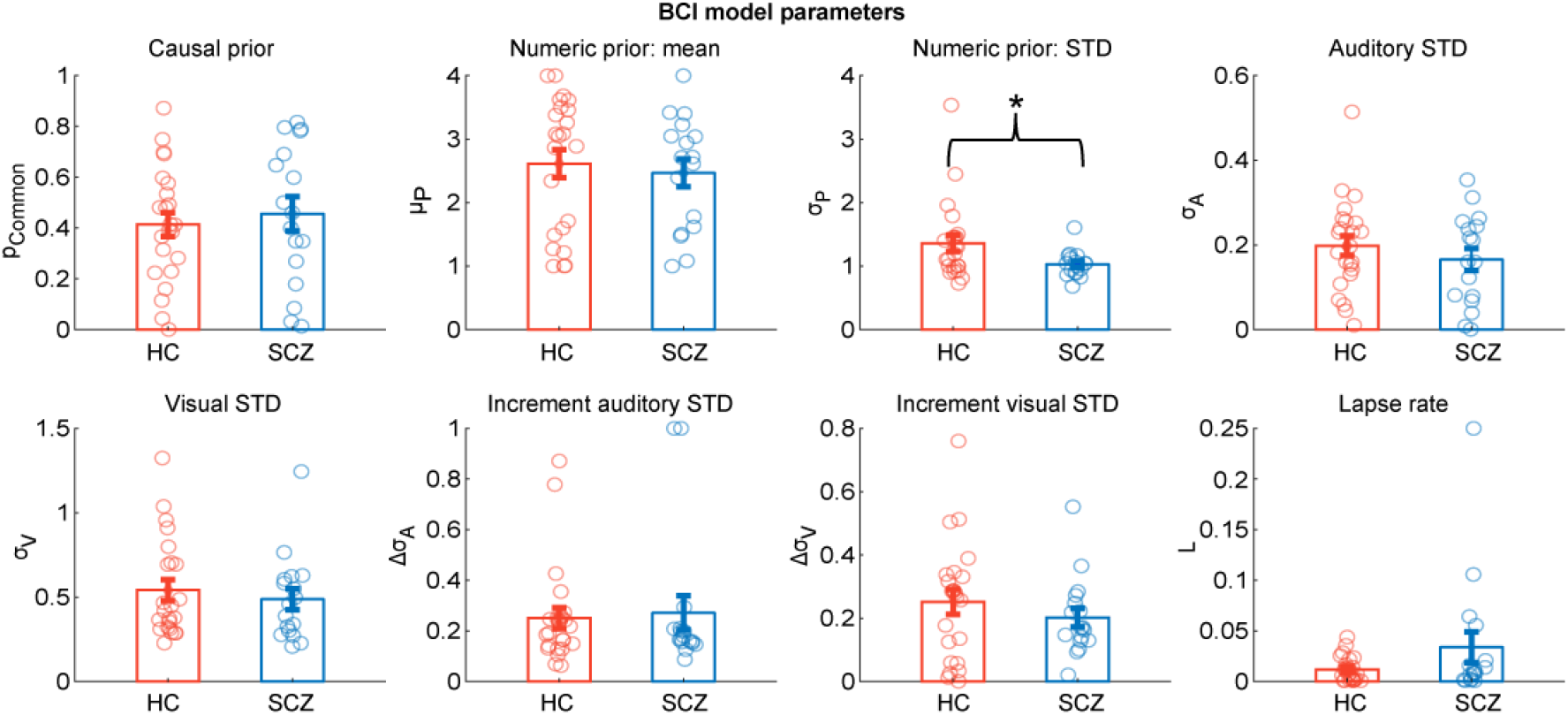
The parameters of the BCI model (across participants mean ± SEM; n = 40) separately plotted for HC and SCZ patients. The BCI model’s decision strategy applies model averaging with increasing sensory variance. Significant different are indicated by * = p < 0.05.

**Table 2.** Comparison of the BCI model’s parameter between HC and SCZ participants.

| | $p_{\text{common}}$ | $\text{var } p_{\text{common}}$ | $\mu_P$ | $\sigma_P$ | $\sigma_A$ | $\sigma_V$ | $\Delta\sigma_A$ | $\Delta\sigma_V$ | $L$ |
| --- | --- | --- | --- | --- | --- | --- | --- | --- | --- |
| $t_{38}$ | -0.534 | 0.764 | 0.454 | 2.109 | 0.938 | 0.599 | -0.275 | 0.958 | -1.658 |
| p | 0.593 | 0.436 | 0.653 | 0.026 | 0.366 | 0.553 | 0.802 | 0.331 | 0.062 |
| Cohen's d | -0.171 | 0.244 | 0.145 | 0.675 | 0.3 | 0.191 | -0.088 | 0.306 | -0.53 |
| $\text{BF}_{10}$ | 0.349 | 0.393 | 0.338 | 1.725 | 0.441 | 0.359 | 0.321 | 0.448 | 0.909 |

Note: The BCI model's decision strategy applies model averaging with increasing sensory variance. p values derived from a two-sample two-sided randomization test ( $n = 5000$ randomizations; uncorrected for multiple comparisons). $p_{\text{common}}$ , causal prior; $\text{var } p_{\text{common}}$ , variance of the causal prior $p_{\text{common}} (1 - p_{\text{common}})$ ; $\mu_P$ , mean of the numeric prior; $\sigma_P$ , standard deviation of the numeric prior; $\sigma_A$ , standard deviation of the auditory likelihood; $\sigma_V$ , standard deviation of the visual likelihood; $\Delta\sigma$ , increment of standard deviation per signal number; $L$ , lapse parameter

In addition, we observed a correlation between negative symptoms and lapse rates in SCZ (r = 0.483, t_16_ = 2.134, p = 0.027, BF_10_ = 1.248) suggesting that these SCZ patients may find it harder to stay attentive throughout the entire experiment. No other significant group differences were revealed when comparing model parameters between the two groups (Tab. 2).

These quantitative Bayesian modelling results corroborate our initial GLM-based conclusions that both HC and SCZ combined audiovisual signals consistent with Bayesian causal inference. The numeric prior was slightly but significantly more precise and the visual variance greater in SCZ compared to HC (according to classical statistics), which is in line with previous results in unisensory perception (Benrimoh et al., 2023; Cassidy et al., 2018; Powers et al., 2017; Teufel et al., 2015).

### Behavior – Adjustment of priors

To further explore whether SCZ over-rely on prior information, we exploited the fact that observers dynamically adapt their priors in response to previous stimuli. Some previous studies have suggested that this dynamic updating of priors is altered in psychosis (Cassidy et al., 2018; but see Valton et al., 2019). Thus, we first examined whether and how SCZ and HC increase their causal prior (i.e. p_common_) after exposure to congruent or incongruent flash-beep sequences. As expected based on previous findings (Gau & Noppeney, 2016; Hong et al., 2022; Nahorna et al., 2012; Rohe et al., 2019), both SCZ and HC similarly increased their causal prior after a congruent trial and decreased it after a trial with large numeric disparity (i.e., a main effect of previous disparity; Fig. 5A and Tab. 3). Importantly, we did not observe any significant differences between groups (i.e. no significant disparity x group interaction).

**Figure 5.**
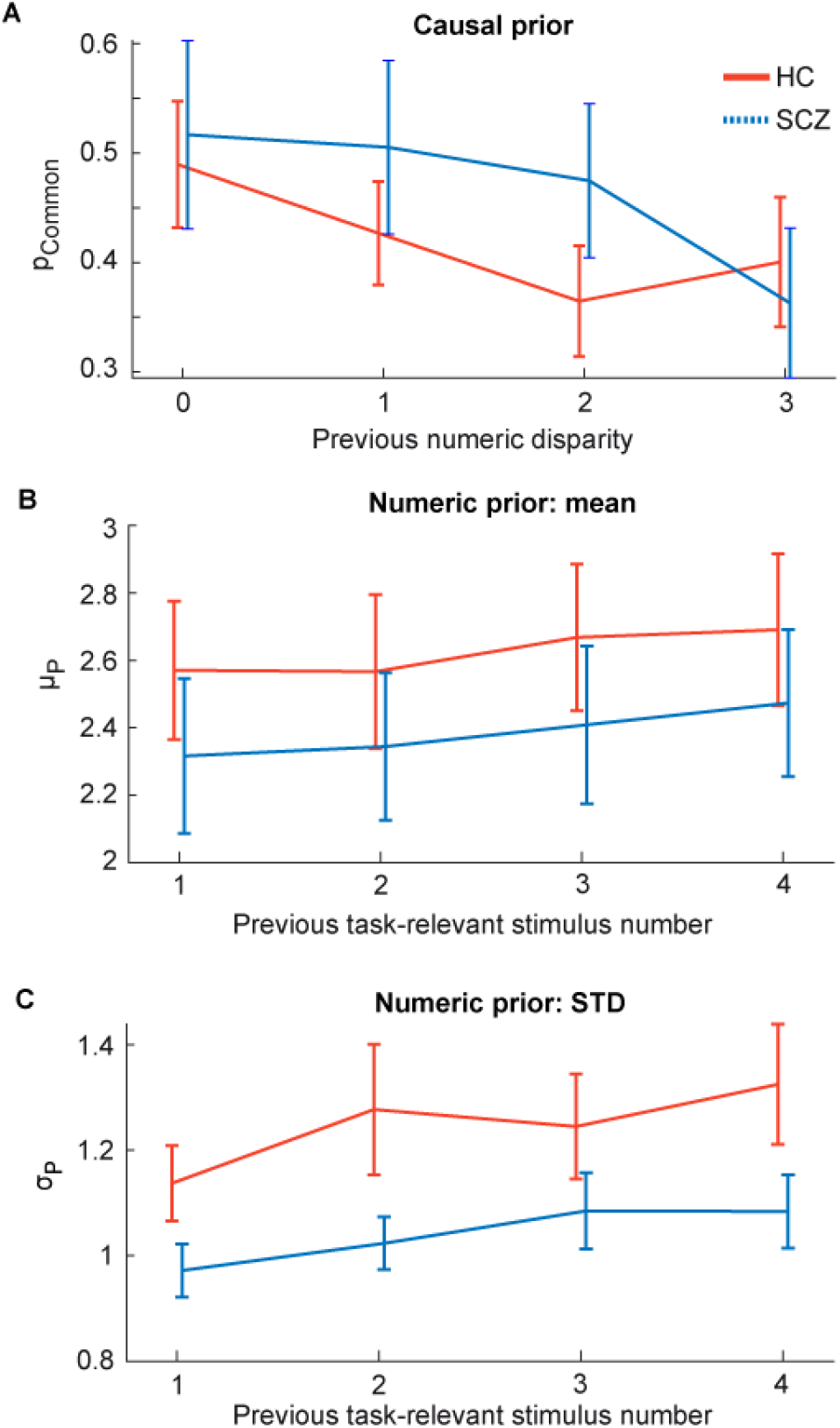
Adjustment of the BCI model’s causal and numeric priors (mean ± between-participants’ SEM; n = 40) in the current trial to the previous trial’s numeric disparity or task-relevant stimulus number in HC and SCZ. **(A)** The current causal prior (p_Common_) as a function of previous numeric disparity (i.e., |n_A_-n_V_|). **(B)** The current numeric prior’s mean (μ_P_) as a function of the previous task-relevant stimulus number (i.e., n_A_ for auditory and n_V_ for visual report). **(C)** The current numeric prior’s STD (σ_P_) as a function of the previous task-relevant stimulus number.

**Table 3.** Main and interaction effects of previous audiovisual disparity (Disp) or audiovisual task-relevant stimulus number (Stim#) and group on the BCI model’s current causal prior (*p*_common_), the numeric prior’s mean (*µ*_P_) and STD (σ_P_) from classical and Bayesian mixed-model ANOVAs.

| Parameter | | F | df1, df2 | p | part. $\eta^2$ | $\text{BF}_{\text{incl}}$ |
| --- | --- | --- | --- | --- | --- | --- |
| $p_{\text{common}}$ | Disp | 5.949 | 3, 114 | <0.001 | 0.135 | 17.903 |
|  | Group | 0.291 | 1, 38 | 0.593 | 0.008 | 0.571 |
|  | Disp×Group | 2.195 | 3, 114 | 0.103 | 0.055 | 0.852 |
| $\mu_P$ | Stim# | 4.820 | 2.6, 97.3 | 0.006 | 0.113 | 7.382 |
|  | Group | 0.568 | 1, 38 | 0.456 | 0.015 | 0.574 |
|  | Stim#×Group | 0.127 | 2.6, 97.3 | 0.923 | 0.003 | 0.130 |
| $\sigma_P$ | Stim# | 6.169 | 3, 114 | <0.001 | 0.140 | 53.296 |
|  | Group | 2.706 | 1, 38 | 0.108 | 0.066 | 0.786 |
|  | Stim#×Group | 0.924 | 3, 114 | 0.432 | 0.024 | 0.361 |

Note: Disp = $|n_A - n_V|$ ; Stim# = $n_A$ or $n_V$ depending on their task relevance; Group: HC vs. SCZ; Effects of the classical mixed-model ANOVAs are Greenhouse-Geisser corrected if sphericity is violated.

Next, we investigated the updating of observers’ numeric prior (i.e. *µ*_P_) and its variance (i.e., σ_P_). As expected for Bayesian learners, both SCZ and HC increased their numeric prior’s mean and variance after exposure to a high number of signals in the task-relevant modality (e.g., beeps for auditory report; Fig. 5B and C; significant main effects in Tab. 3). But again, no significant differences were observed between groups (i.e., no significant interaction effects with group) suggesting that HC and SCZ also adjust their perceptual priors at shorter timescales similarly.

In summary, our analyses of the behavioral data based on the general linear model (GLM) and formal Bayesian modelling suggest that the medicated SCZ dynamically adapted and combined priors about the signals’ causal structure and the number of signals with new audiovisual inputs comparable to the healthy controls and in line with Bayesian principles.

### EEG – Decoding of numeric Bayesian Causal Inference estimates

To investigate whether SCZ and HC achieve Bayesian causal inference via shared or different neural processes, we combined BCI models with EEG decoding. In particular, we temporally resolved how the brain encodes the auditory (*N̂*_A,C=2_) and visual (*N̂*_V,C=2_) segregation, the fusion (*N̂*_AV,C=1_) and the final BCI perceptual estimates (i.e. *N̂*_A_ or *N̂*_V_) that combine the forced-fusion estimate with the task-relevant unisensory segregation estimates, weighted by the posterior causal probability of a common or separate causes (Rohe et al., 2019). To track the evolution of these different estimates across time, we trained a linear support-vector regression model (SVR decoder) to decode each of the four BCI estimates from EEG patterns of 60 ms time windows. In both HC and SCZ, the decoders predicted the BCI estimates from EEG patterns significantly better than chance over most periods of the post-stimulus time window (Fig. 6A). Yet, the different BCI estimates evolved with different time courses. Initially, the EEG activity encoded mainly the visual segregation estimates starting at about ∼60-100 ms (i.e., significant clusters in one-sided cluster-based corrected randomization t-test; see Supporting Tab. S4). Slightly later, the auditory segregation and fusion estimates peaked with the fusion estimates showing a slower decline. The final BCI estimate, which accounts for the signals’ causal structure, rose more slowly and showed a more sustained time course. Moreover, in HC only did its decoding accuracy exceed those of the other estimates at 600 ms. The temporal profile of the decoding accuracies were largely comparable between HC and SCZ. Bayes factors provided mainly weak evidence for no difference between the groups (Fig. 6B), except for the visual segregation-estimate that was associated with a significantly lower decoding accuracyin SCZ than HC from 120 to 220 ms (Fig. 6A and Supplemental Tab. S4; all further clusters p > 0.05). Because we did not track observers’ eye movements, these lower decoding accuracies in SCZ may potentially be explained by reduced fixation stability in SCZ (Benson et al., 2012). In summary, combining Bayesian modelling and EEG decoding largely confirmed that SCZ and HC perform audiovisual number estimation according to similar neurocomputational mechanisms. However, SCZ encoded the visual signal with less precision resulting in a lower decoding accuracy of the visual segregation estimate.

**Figure 6.**
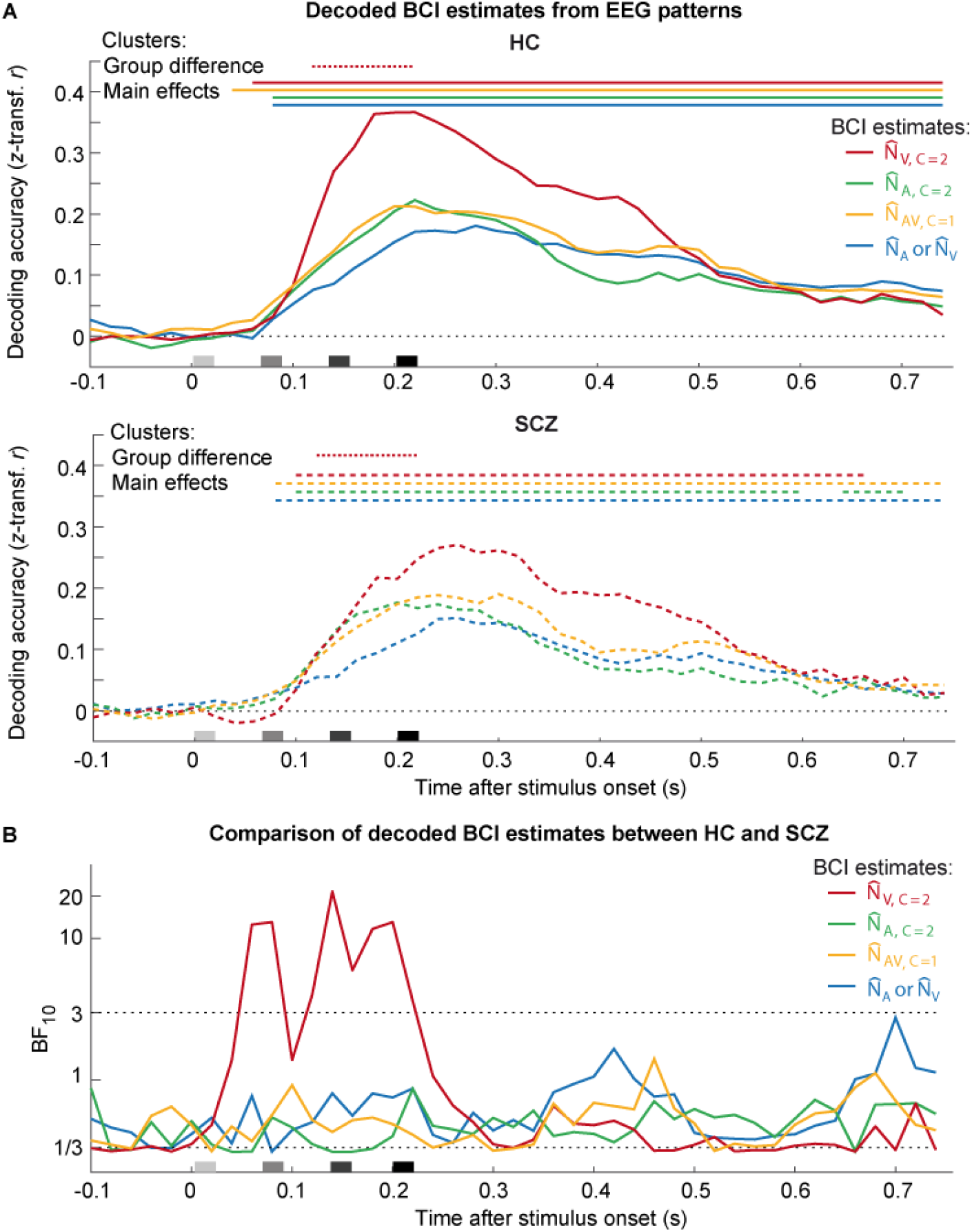
Decoding the BCI model’s numeric estimates from EEG patterns using support-vector regression (SVR) in HC versus SCZ (n = 40). **(A)** Decoding accuracy (Fisher’s *z*-transformed correlation; across-participants mean) of the SVR decoders as a function of time and group (HC vs. SCZ). Decoding accuracy was computed as Pearson correlation coefficient between the given BCI model’s internal estimates and BCI estimates that were decoded from EEG activity patterns using SVR models trained separately for each numeric estimate. The BCI model’s internal numeric estimates comprise of: i. the unisensory visual (*N̂*_V,C=2_), ii. the unisensory auditory (*N̂*_A,C=2_) estimates under the assumption of independent causes (*C*=2), iii. the forced-fusion estimate (*N̂*_AV,C=1_) under the assumption of a common cause (*C*=1) and iv. the final BCI estimate (*N̂*_A_ or *N̂*_V_ depending on the sensory modality that is task-relevant) that averages the task-relevant unisensory and the precision-weighted estimate by the posterior probability estimate of each causal structure. Color-coded horizontal solid lines (HC) or dashed lines (SCZ) indicate clusters of significant decoding accuracy (*p* < 0.05; one-sided one-sample cluster-based corrected randomization t-test). Color-coded horizontal dotted lines indicate clusters of significant differences of decoding accuracy between both groups (*p* < 0.05; two-sided two-sample cluster-based corrected randomization t-test). Stimulus onsets are shown along the x-axis. **(B)** Bayes factors for the comparison between the decoding accuracies of HC and SCZ for each of the BCI estimate (i.e., BF_10_ > 3 substantial evidence for or BF_10_ < 1/3 against group differences).

### EEG – Multisensory interactions in ERPs

Following previous work, we also analyzed and compared the event related potentials (ERP) to unisensory and multisensory stimuli between HC and SCZ (Balz et al., 2016; Roa Romero, Keil, Balz, Gallinat, et al., 2016; Roa Romero, Keil, Balz, Niedeggen, et al., 2016; Stekelenburg et al., 2013; Stone et al., 2014; Stone et al., 2011). ERPs showed the typical components in response to flashes and beeps (Fig. 7), i.e. P1 (∼ 50 ms), N1 (100 ms), P2 (200 ms), N2 (280 ms) and P3 (> 300 ms) (Mishra et al., 2007). As previously reported (Oribe et al., 2015), the unisensory visual P3 component was significantly smaller in SCZ than HC (cluster 560*–*705 ms, p = 0.045). To characterize the neural processes of audiovisual integration, we tested for audiovisual interactions (i.e., AV_congr_ vs. (A + V)) over occipital electrodes. In line with previous studies (Mishra et al., 2007; Shams et al., 2005), we observed early audiovisual interactions from 75*–*130 ms (i.e., measured from the onset of the first flash-beep slot; *p* = 0.049; two-sided one-sample cluster-based corrected randomization test) and later negative audiovisual interactions from 260-750 ms after stimulus onset (p < 0.001). Crucially, the ERP interactions did not differ between HC and SCZ (p > 0.05; two-sided two-sample cluster-based corrected randomization test).

**Figure 7.**
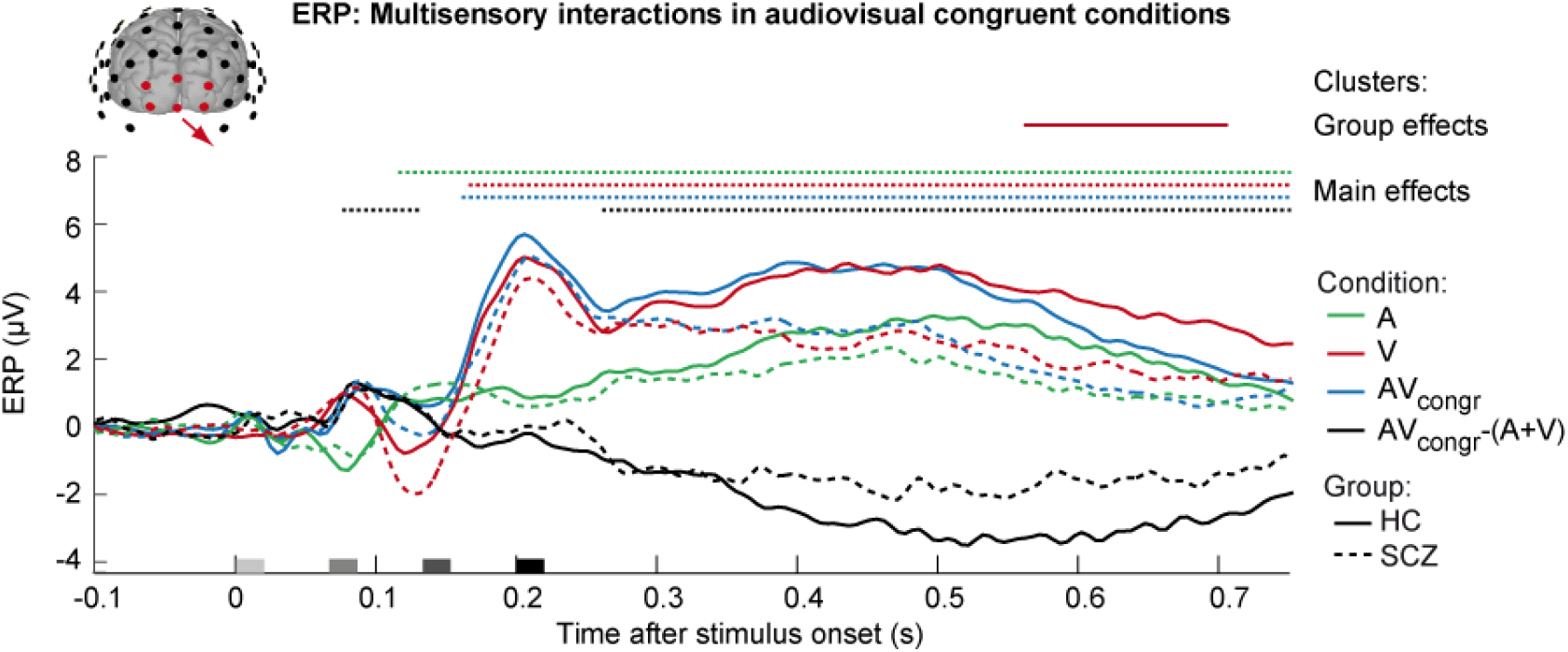
Occipital ERPs in response to unisensory and audiovisual stimuli in HC and SCZ. Event-related potentials (ERPs; across-participants mean grand averages; n = 40) of HC and SCZ participants elicited by unisensory auditory stimuli (A), unisensory visual stimuli (V), audiovisual congruent conditions (AV_congr_) and the difference of these ERPs (i.e., AV_congr_ – (A+V)) indicating multisensory interactions. The ERPs are averaged across occipital electrodes. Color-coded horizontal dotted lines indicate significant clusters (p < 0.05) of ERPs against baseline (i.e., across HC and SCZ, main effect of condition) in one-sample two-sided cluster-based corrected randomization tests. The horizontal solid line indicates a significant cluster of ERP differences between HC and SCZ in two-sample two-sided cluster-based corrected randomization tests. The x-axis shows the stimulus onsets.

To examine whether SCZ patients and HC may differ more subtly in their multivariate EEG response, we trained a support-vector classifier (SVC) to classify participants (Bae et al., 2020) as SCZ or HC based on their EEG activation patterns to auditory, visual and audiovisual stimuli across poststimulus time windows (see Supporting methods and Fig. S4). The ERP topographies evolved remarkably similar in HC and SCZ, so that the EEG-based decoder did not classify participants as HC or SCZ better than chance in one-sided cluster-based corrected randomization tests when corrected across multiple comparisons across time. Overall, our ERP and multivariate analyses thus corroborated that the neural mechanisms underlying audiovisual integration were largely comparable in HC and SCZ patients. However, in line with our EEG results from the BCI analysis we observed a deficit in visual processing in SCZ as indicated by smaller visual P3 responses.

## Discussion

This study combined psychophysics, EEG and Bayesian modelling to investigate whether and how schizophrenia impacts the computational and neural mechanisms of multisensory causal and perceptual inference. A growing number of studies suggests that schizophrenia may alter the brain’s ability to integrate audiovisual signals into coherent percepts (for a review, see Tseng et al. (2015)). Many of these studies have used qualitative approaches such as evaluating the frequency of perceptual illusions in schizophrenia compared to healthy controls. Yet, the complex dependencies of multisensory illusion rates on sensory precisions and perceptual as well as causal priors may have contributed to the inconsistent findings reported so far (Balz et al., 2016; Haß et al., 2017; Vanes et al., 2016). In this study, we have therefore employed a more sophisticated factorial design with formal Bayesian modelling to disentangle these different computational components as has been successfully shown for autism spectrum disorder (Noel et al., 2020; Noel et al., 2022).

Our GLM-based and Bayesian modelling analyses offer convergent evidence that SCZ as their HC counterparts perform multisensory perceptual and causal inference in line with normative principles of Bayesian causal inference. When numeric disparities are small and signals are likely to originate from common sources, individuals with SCZ and HC integrate auditory and visual signals weighted by their relative precisions resulting in crossmodal biases. At large numeric disparities, these crossmodal biases and interactions are reduced. Through formal Bayesian model comparison, we further examined whether SCZ compute Bayesian estimates or resort to simpler heuristics such as segregating signals above a numeric disparity threshold. Our quantitative Bayesian modelling analyses confirmed that SCZ, like their healthy counterparts, combine signals consistent with BCI models and read out final numeric estimates based on the decisional strategy of model averaging.

Next, we examined whether SCZ may overweight prior information about flash/beep number in relation to sensory evidence at long and/or short timescales as previously observed in unisensory perception (Benrimoh et al., 2023; Cassidy et al., 2018; Powers et al., 2017; Teufel et al., 2015). In support of an overweighting of priors in psychosis, our Bayesian modelling analysis revealed a significantly more precise numeric prior for SCZ compared to HC. Further, the visual variance correlated with positive symptoms on the PANSS scale. These findings are consistent with previous studies showing abnormal weighting of prior knowledge and sensory evidence in SCZ. However, we cannot exclude the possibility that SCZ may have been less vigilant and failed to count some signals, which could be misinterpreted as overreliance on their numeric prior (see Fig. 4, 5B). In support of intact weighting of prior and sensory information at shorter timescales, the influence of the previous trial’s flash/beep number on current perceptual choices was comparable in SCZ and HC. Overall, our analyses thus suggest that the medicated SCZ group of our study weighted and dynamically adapted numeric priors largely comparable to their healthy counterparts with a small trend towards a more precise sustained numeric prior.

Our multisensory paradigm enabled us to assess not only the weighting of prior knowledge regarding environmental properties such as flash/beep number, but also the world’s causal structure, i.e. whether signals come from common or independent sources as incorporated in the causal prior. A high causal prior or binding tendency increases crossmodal biases and interactions, while a low causal prior enhances an individual’s ability to selectively report the signal number in the task-relevant sensory modality whilst ignoring the incongruent number of signals in the irrelevant sensory modality. Changes in the causal prior may thus be closely associated with conflict monitoring, cognitive control and selective attention mechanisms (Badde et al., 2020; Ferrari & Noppeney, 2021; Odegaard et al., 2016; Rohe & Noppeney, 2018). Yet, despite growing evidence for impaired selective attention and cognitive control mechanisms in SCZ (Gold et al., 2007; Heinrichs & Zakzanis, 1998; Lesh et al., 2011), our Bayesian modelling analyses did not reveal any significant differences in the causal prior between SCZ and HC.

In summary, our GLM-based and Bayesian modelling analyses showed that SCZ and HC combined perceptual and causal priors with sensory evidence in a manner consistent with the principles of normative Bayesian causal inference. Furthermore, Bayesian statistics provided consistent evidence that the causal and perceptual priors’ weight and updating was largely maintained in the SCZ group. This absence of notable computational abnormalities could be explained by the fact that our patient group was post-acute and medicated and scored low on the PANSS positive symptom scale. Most importantly, our cohort of SCZ patients also showed near-normal performance on the TMT-B and Stroop tests as indices of executive and attentional functions which are particularly relevant for optimal performance in our inter-sensory selective attention paradigm (cf. supporting Tab. S1). Our findings add further evidence to the fact that fundamental mechanisms of perceptual inference are preserved at least in some medicated SCZ individuals (Jardri et al., 2017; Schmack et al., 2017; Valton et al., 2019; Weilnhammer et al., 2020). Future research is required to explore whether schizophrenia patients with attentional and executive deficits may exhibit abnormal causal priors, and thus, how they arbitrate between sensory integration and segregation.

Using EEG, we investigated whether SCZ and HC support these computations via similar or different neural mechanisms. For instance, in SCZ, the computations may involve compensatory neural systems or exhibit a slower dynamics. To explore these questions, we decoded the auditory and visual segregation, the fusion and the final BCI model’s estimates from scalp EEG data. In HC, the decoding accuracy of the visual segregation-estimate peaked earlier and higher compared to the other estimates. By contrast, in the SCZ group, decoding of the visual estimate showed a more protracted time course initially overlapping with the auditory segregation and fusion estimates. Statistical tests confirmed a significant difference in decoding accuracy between HC than SCZ from approximately 100 – 200 ms. However, apart from this difference in decoding of the unisensory visual estimate and similarly in the visual P3 ERP component, the decoding profiles appeared remarkably similar across the two groups.

To ensure that we did not miss any differences between SCZ and HC that were previously reported in the literature (Balz et al., 2016; Roa Romero, Keil, Balz, Gallinat, et al., 2016; Roa Romero, Keil, Balz, Niedeggen, et al., 2016; Stekelenburg et al., 2013; Stone et al., 2014; Stone et al., 2011), we also performed standard ERP analyses to test for audiovisual interactions. However, these analyses revealed only an attenuated visual P3 component in SCZ, but no significant differences in audiovisual interactions between the two groups. Similarly, we were unable to predict group membership (i.e., HC vs. SCZ) from multivariate audiovisual EEG responses significantly better than chance.

To conclude, our behavioral, computational and neuroimaging results consistently demonstrate that audiovisual perception in a sound-induced flash illusion paradigm is based on comparable computational and neural mechanisms in SCZ and HC. Both SCZ and HC combined audiovisual signals into number estimates in line with the computational principles of Bayesian causal inference. Further, time-resolved EEG decoding revealed that both HC and SCZ performed Bayesian causal inference by dynamically encoding perceptual estimates that segregate, integrate and flexibly combine auditory and visual information according to their causal structure. Our results show that at least in our group of post-acute medicated SCZ patients, the computations and neural mechanisms of hierarchical Bayesian causal inference in audiovisual perception were remarkably intact. Future research is needed to determine whether deviations from normative Bayesian principles in multisensory perception may occur in unmedicated patients in prodromal or acute stages or patient groups with more pronounced impairments on their attentional and executive functions. Critically, our study focused on simple artificial flash-beep stimuli. This makes it an important future research direction to investigate multisensory perceptual inference in psychosis in more naturalistic situations such as face-to-face communication.

## Materials and Methods

### Participants

After giving written informed consent, 23 healthy volunteers and 17 post-acute in- and out-patients with schizophrenia were included in the study. Healthy volunteers and individual with schizophrenia were matched in age, sex and education (Supporting Tab. S1; cf. the supporting information for further details on the sample and clinical experimental procedures). Data from the 23 healthy participants were reported in Rohe et al. (2019). The study was approved by the human research review committee of the Medical Faculty of the University of Tuebingen and at the University Hospital Tuebingen (approval number 728/2014BO2).

### Experimental design

In the flash-beep paradigm, participants were presented with a sequence of i. one, two, three or four flashes and ii. one, two, three or four beeps (Fig. 1A; cf. supporting information for further details on the stimuli and the experimental setup). On each trial, the number of flashes and beeps was independently sampled from one to four leading to four levels of numeric audiovisual disparities (i.e. zero = congruent to four = maximal level of disparity; Fig. 1B). Each flash and/or beep was presented sequentially in fixed temporal slots that started at 0, 66.7, 133, 200 ms. The temporal slots were filled up sequentially. For instance, if the number of beeps was three, they were presented at 0, 66.6, 133 and 200 ms, while the fourth slot was left empty. Hence, if the same number of flashes and beeps was presented on a particular trial, beeps and flashes were presented in synchrony. On numerically disparate trials, the ‘surplus’ beeps (or flashes) were added in the subsequent fixed time slots (e.g. in case of 2 flashes and 3 beeps: we present 2 flash-beeps at 0 and 66.6 ms in synchrony and a single beep at 133 ms).

Across experimental runs, we instructed participants to selectively report either the number of flashes or beeps and to ignore the stimuli in the task-irrelevant modality. Hence, the 4 × 4 × 2 factorial design manipulated (i) the number of visual flashes (i.e. one, two, three or four), (ii) the number of auditory beeps (i.e. one, two, three or four) and (iii) the task relevance (auditory-vs. visual-selective report) yielding 32 conditions in total (Fig. 1B). For analyses of the crossmodal bias, we reorganized trials based on their absolute numeric disparity (|#A - #V| ∈ {0,1,2,3}).

### Overview of GLM and Bayesian modelling analyses for behavioral data

We assessed similarities and differences in perceptual and causal inference between SCZ and HC by combining general linear model (GLM)-based and a Bayesian modelling analysis approaches. The GLM-based analysis computed i. the correlation between true and reported number of flashes/beep for congruent and unisensory trials and ii. the crossmodal bias (CMB) which quantified the relative influence of the auditory and the visual numeric stimuli on observers’ auditory and visual behavioral numeric reports. The Bayesian modelling analysis fitted BCI models with different decisional strategies and additional heuristic models to the behavioral numeric reports. We then used Bayesian model comparison to determine the model(s) that were the best explanation for the behavioral data in SCZ and/or HC.

### GLM-based analysis of behavioral data

Our GLM-based analysis of behavioural data focused on response accuracy indices for unisensory and congruent conditions and on the crossmmodal bias for incongruent audiovisual conditions. To compare overall task performance between HC and SCZ, we computed participants’ response accuracy by correlating participants’ numeric report with the true task-relevant number of auditory or visual stimuli (i.e., Pearson correlation coefficient). We then analyzed Fisher z-transformed correlation coefficients in unisensory and audiovisual congruent conditions using a mixed-model ANOVA with within-participants factors modality (unisensory vs. audiovisual) and task relevance (auditory vs. visual report) and between-participant factor group (HC vs. SCZ) (Fig. 1C and Tab. 1).

We characterized how HC and SCZ participants weight audiovisual signals during multisensory perception by computing the crossmodal bias (CMB) from their responses to audiovisual incongruent stimuli (Rohe & Noppeney, 2015b; Wallace et al., 2004). The CMB quantifies the relative influence of the auditory (n_A_) and the visual (n_V_) numeric stimuli on observers’ auditory and visual behavioral numeric reports (r_A/V_):

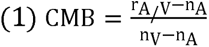

To account for response biases, we adjusted n_A_ and n_V_ with a linear regression approach across all congruent trials in a participant-specific fashion. In other words, we replaced the true n_A_ and n_V_ in the CMB equations with the n_A_ and n_V_ predicted based on participants’ responses during the congruent conditions (Rohe & Noppeney, 2015b). This CMB-based analysis allowed us to compare the audiovisual weight profiles between HC and SCZ using a mixed-model ANOVA with within-participant factors numeric disparity (1, 2, 3) and task relevance (auditory vs. visual report) and between-participant factor group (HC vs. SCZ) (Fig. 1D and Tab. 1). Further, the analysis allowed us to assess whether the weight profiles were qualitatively in line with the principles of Bayesian causal inference (i.e. a strong crossmodal bias for smaller relative to larger numeric disparities). Complementary Bayesian ANOVAs were used to provide evidence for no difference between the groups (see section: ‘Statistical analysis of behavioural data’ below).

### Bayesian modelling analysis for behavioral data

The Bayesian modelling analysis fitted several competing computational models to the individual behavioral numeric reports. We then used factorial Bayesian model comparison to determine the model that is the best explanation for observers’ behavioral data and compared the results of the model comparison and model parameters between HC and SCZ (Tab. 2).

In the following, we will briefly describe the competing models which spanned a two-factorial model space. The first factor compared five model types that differed in their decision function, i.e. how they combined the estimates of the fusion and the segregation components into a final decision. The three BCI model combined the fusion and segregation estimates according to the decisional strategies of model averaging, model selection and probability matching (Wozny et al., 2010). Two additional heuristic models reported the fusion or segregation estimates with a fixed probability (stochastic fusion) or depending on whether the numeric disparity was greater than a fixed threshold (i.e., fixed-criterion threshold model; (Acerbi et al., 2018)). The second factor compared whether auditory and visual variances were equal across all number of beeps/flashes or increased with stimulus number (i.e. scalar-variability model; (Dehaene, 2007; Gallistel & Gelman, 2000)). Details on the BCI model can be found in Kording et al. (2007) and Acerbi et al. (2018). Specific details on fitting the BCI model to numeric reports in the current experimental paradigm can be found in the supporting information and in Rohe et al. (2019).

### Statistical analyses of behavioral data

To refrain from making parametric assumptions, we compared behavioural measures and parameters (i.e., neuropsychological test scores, CMB and BCI model parameters; Tab. S1 and Tab. 2) between conditions and groups as well as tested correlations between BCI model parameters and symptom scores using randomization tests with t-values as test statistics (n = 5000 randomizations). To assess the evidence in favor of the null-hypothesis (i.e. no difference between groups or conditions), randomization tests were complemented with Bayesian analyses.

To evaluate whether experimental factors influenced HC and SCZ differentially or equivalently (e.g., for CMB), we complemented classical with Bayesian mixed-model ANOVAs (Tab. 1 and 3). For Bayesian analyses, the following interpretations of BF apply: BF > 3 or > 10 provide substantial or strong evidence (Kass & Raftery, 1995; Wagenmakers et al., 2018) for condition/group differences, correlations or inclusion of a factor, whereas BF < 1/3 or < 1/10) provides substantial or strong evidence for condition/group equivalence, no correlation or exclusion the factor (van den Bergh et al., 2020). For further details on the statistical analysis, please note the supporting information.

### EEG – Data acquisition and preprocessing

EEG signals were recorded from 64 active electrodes positioned in an extended 10–20 montage using electrode caps (actiCap, Brain Products, Gilching, Germany) and two 32 channel DC amplifiers (BrainAmp, Brain Products). For analysis of event-related potentials (ERPs) and decoding analyses (see below), all EEG data were baseline corrected with a 200 ms prestimulus baseline and were analyzed from 100 ms before stimulus onset up to 750 ms after stimulus onset, when the response screen was presented (cf. supporting information for futher details EEG data acquisition and preprocessing).

For multivariate analyses (see below), single-trial EEG data from the 64 electrodes were binned in sliding time windows of 60 ms using an overlap of 40 ms. Hence, given a sampling rate of 200 Hz, each 60 ms time window included 12 temporal sampling points. 64-electrode EEG activity vectors (for each time sample) were concatenated across the 12 sampling points within each bin resulting in a spatiotemporal EEG activity pattern of 768 features. EEG activity patterns were z scored to control for mean differences between conditions. The first sampling point in the 60 ms time window was taken as the window’s time point in all analyses.

### EEG – Analysis of audiovisual interactions in ERPs

In univariate analyses, we assessed whether schizophrenia alters basic sensory components and early audiovisual interactions in occipital ERPs. We averaged trial-wise EEG data time-logged to stimulus onset into ERPs for audiovisual congruent conditions (i.e., controlling for effects of disparity and averaging over effects of task-relevance) and unisensory conditions. We then averaged the ERPs across occipital electrodes (i.e. O1, O2, Oz, PO3, POz, PO4; Fig. 7). To analyze early audiovisual interactions, we computed the difference between congruent audiovisual conditions and the corresponding unisensory conditions (i.e., AVcongr – (A + V)). Because attentional and cognitive sets between uni- and bisensory runs might have differed and our experimental design did not include null trials to account for anticipatory effects around stimulus onset (Teder-Sälejärvi et al., 2002), the audiovisual interactions need to be interpreted with caution. To test whether the difference waves deviated from zero at the group level across HC and SCZ, we used a non-parametric randomization test (5000 randomizations) in which we flipped the sign of the individual difference waves using a one-sample t-test as a test statistic (Nichols & Holmes, 2002). To compare the difference waves between both groups, we used a randomization test in which we flipped group membership (5000 randomizations) using two-sample t-tests as a test statistic. To correct for multiple comparisons across the EEG sampling points, we used a cluster-based correction (Maris & Oostenveld, 2007) with the sum of the t values across a cluster as cluster-level statistic and an auxiliary cluster-defining threshold of t = 2 for each time point. In a supplementary analysis, we used a group-classification approach (Bae et al., 2020), in which we assessed whether the multivariate EEG response patterns to unisensory and multisensory stimuli differed between HC and SCZ (Supporting methods and results; Fig. S4).

### EEG – Multivariate analyses of BCI estimates

To characterize whether the neurophysiological processes underlying Bayesian causal inference differ between HC and SCZ, we decoded the four numerical estimates of the BCI model from multivariate EEG patterns across poststimulus time points using linear support-vector regression (Rohe et al., 2019; Rohe & Noppeney, 2015a) as implemented in LibSVM 3.20 (Chang & Lin, 2011). The individually fitted BCI model provided the i. unisensory visual (*N̂*_V,C=2_), ii. unisensory auditory (*N̂*_A,C=2_) estimates, iii. forced-fusion (*N̂*_AV,C=1_), iv. final BCI audiovisual numeric estimate (*N̂*_A_ or *N̂*_V_) for each of the 32 audiovisual conditions (see above). For each numeric estimate and each time window, an SVR model was trained to decode the numeric estimate from single-trial EEG activity patterns (see above for a definition) across all 32 conditions in a leave-one-run-out cross validation scheme. The SVRs’ parameters (C and *v*) were optimized using a grid search within each cross-validation fold (i.e., nested cross-validation).

To quantify the decoding accuracies, we computed the correlation coefficients between the ‘true’ model estimates and the decoded model estimates (Fig. 6A). The Fisher’s z-transformed correlation coefficients were tested against zero across HC and SCZ using a one-sided randomization test (i.e., sign flip of correlation coefficient in 5000 randomizations; one sample t-test as test statistic). The Fisher’s z-transformed correlation coefficients were compared between HC and SCZ using a two-sided randomization test (i.e., 5000 randomizations; two-sample t-tests as a test statistic). Further, we applied Bayesian two-sample t-tests (as described above for behavioral data) to compute Bayes factors that quantify evidence for or against a group differences in decoding accuracies across poststimulus time points (Fig. 6B).

## Supporting information

Supplemental methods and results

## Acknowledgements

This study was funded by the University of Tuebingen (Fortüne grant numbers 2292–0–0 and 2454–0–0) and the Deutsche Forschungsgemeinschaft (DFG; grant number RO 5587/ 1–1). We thank Luigi Acerbi for sharing code to perform robust Bayesian model-selection procedures which was updated by David Meijer. We thank Ramona Täglich and Larissa Metzler for help with the data collection.

