## Supplemental methods and results for "Computations and neural dynamics of audiovisual causal and perceptual inference in schizophrenia"

**Supporting Information Text**

*Participants*

After giving written informed consent, 24 healthy volunteers and 26 post-acute in- and out-patients with psychosis participated in the EEG study based on previous calculations of statistical power. The calculations yielded sufficient statistical power (~0.97) assuming a large effect size (i.e, Cohen’s d = 1.00) from older previous studies showing reduced multisensory integration in schizophrenia patients (de Gelder et al., 2003; Pearl et al., 2009; Williams et al., 2010). Healthy participants were included when meeting the inclusion criteria (age 20-65, adequate German comprehension, normal or corrected-to-normal vision and audition) and not the exclusion criteria (i.e., no psychiatric or psychosomatic disorders except specific phobias and nicotine addiction; no cardio-vascular disorders, diabetes and neurological disorders). Healthy participants were matched to patients for age, sex and education (Tab. S1). Current or former psychiatric disorders were screened using the questions of the structured clinical interview for DSM IV axis I disorders, SCID-I, German version. Patients were included when meeting the same inclusion- and exclusion criteria as healthy controls (except psychiatric disorders). To clinically diagnose the patients, they underwent the full SCID-I interview (see below). 19 of the 26 patients fulfilled the criteria for schizophrenia and were included (12 paranoid, 1 disorganized, 3 undifferentiated, 3 residual). 6 patients fulfilled the criteria for schizoaffective disorder and 1 patient for substance-induced psychotic disorder and were excluded (note that including these patients leads to similar results). Two of the 19 SCZ patients were excluded because they showed an unusual response profile in the CMB with a very small, outlying task effect (i.e., task effect < 2 STD from the mean task effect) suggesting that these two patients only used the beeps to provide the auditory and visual numeric reports.

Of the 17 included SCZ patients, 4 exhibited comorbid major depression (lifetime), one patient alcohol dependence (lifetime), three patients cannabis dependence (lifetime, one also acute), one patient cocaine dependence (lifetime), one patient social phobia (lifetime and acute), two patients compulsive-obsessive disorder (lifetime), one patient binge-eating disorder (acute; for general demographics and other sample characteristics, see Tab. S1). The mean duration of the psychotic illness in SCZ patients was 13.37 ± 10.75 years (mean ± STD). 15 patients were medicated with atypical antipsychotics (amisulpride, aripriprazole, quetiapine, olanzapine, paliperidone), 1 patient was medicated with typical and atypical antipsychotics (haloperidol and clozapine) and 1 patient was unmedicated. Acute positive symptoms were rather moderate in the current sample (cf. PANSS, Tab. S1).

*Experimental procedures*

Participants took part in two sessions on two separate days. In the first session, participants underwent clinical and neuropsychological assessments (Tab. S1). For HC, clinical assessments comprised the screening of the SCID-I interview (Wittchen et al., 1997). For SCZ, clinical assessment comprised the full SCID-I interview, assessment of the Positive and Negative Symptom Scale (PANSS)(Kay et al., 1987), the Calgary Depression Scale for Schizophrenia (CDSS) (Addington et al., 1990) and a recording of clinical data (e.g., medication). Antipsychotic medication was converted to chlorpromazine-equivalent doses (Leucht et al., 2014). Both groups underwent neuropsychological tests comprising the verbal learning and memory test (VLMT) (Helmstaedter et al., 2001), the Trail Making Test (TMT-A and -B) (Reitan, 1992) assessing visual attention and executive functions (i.e., task switching), a test for premorbid crystallized intelligence (Mehrfachwahl-Wortschatz-Intelligenztest, MWT-B) (Lehrl, 2005), a Stroop test (Bäumler, 1984) assessing executive functions (i.e., cognitive inhibition), the Edinburgh Handedness Inventory (EHI) (Oldfield, 1971) and Beck’s Depression Inventory (BDI) (Hautzinger et al., 2006). All clinical and neuropsychological assessment were made by trained and experienced clinicians. In the second session, participants underwent a flash-beep paradigm including EEG measurements.

*Stimuli*

The flash-beep paradigm was an adaptation of the “sound-induced flash illusion” paradigm (Shams et al., 2000; Shams et al., 2005). The visual flash was a circle presented in the center of the screen on a black background (i.e. 100% contrast; Fig. 1A) briefly for one frame (i.e, 16.7 ms, as defined by the monitor refresh rate of 60 Hz). The maximum grayscale value (i.e. white) of the circle was at radius 4.5° with smoothed inner and outer borders by defining the grayscale values of circles of smaller and larger radius by a Gaussian of 0.9° STD visual angle. The auditory beep was a pure tone (2000 Hz; ~ 70 dBSPL) of 27 ms duration including a 10 ms linear on/off ramp. Multiple visual flashes and auditory beeps were presented sequentially at a fixed SOA of 66.6 ms.

On each trial, the number of flashes and beeps was independently sampled from one to four leading to four levels of numeric audiovisual disparities (i.e. zero = congruent to four = maximal level of disparity; Fig. 1B). Each flash and/or beep was presented sequentially in fixed temporal slots that started at 0, 66.7, 133, 200 ms. The temporal slots were filled up sequentially. The duration of a flash-beep sequence was determined by the number of sequentially presented flash and/or beep stimuli. Irrespective of the number of flashes and/or beeps, a response screen was presented 750 ms after the onset of the first flash and beep for a maximum duration of 2.5 s instructing participants to report their perceived number of flashes (or beeps) as accurately as possible by pushing one of four buttons. The order of buttons was counterbalanced across runs to decorrelate motor responses from numeric reports. On half of the runs, the buttons from left to right corresponded to one to four stimuli; on the other half, they corresponded to four to one. After a participant’s response, the next trial started after an inter-trial interval of 1–1.75 s.

In every experimental run, each of the 16 conditions (i.e., 1-4 visual flashes x 1-4 auditory beeps) was presented 10 times. Healthy control participants completed 4 runs of auditory- and 4 runs of visual-selective report in a counterbalanced fashion (except for one participant performing 5 runs of auditory and 3 of visual report) and two unisensory runs with visual or auditory stimuli only (i.e. 4 unisensory conditions presented 40 times per run). Thus, HC participants completed 10 runs in total. SCZ participants completed 4-8 runs of auditory- and visual-selective report, depending on their endurance, and one unisensory visual and one auditory run. Thus, SCZ participants completed 6-10 runs in total (7.47 ± 0.36, mean ± SEM number of runs). To control for time-dependent differences between groups, the trial numbers of behavioral and EEG data for each HC individual were subsampled to match the average trial number of the SCZ group across all analyses (i.e., trials 1-1186 were taken in HC). Before the actual experiment, participants completed 56 practice trials.

*Experimental setup*

Psychtoolbox 3.09 (Brainard, 1997) (www.psychtoolbox.org) running under MATLAB R2016a (MathWorks) presented audiovisual stimuli and sent trigger pulses to the EEG recording system. Auditory stimuli were presented at ≈ 70 dB SPL via two loudspeakers (Logitech Z130) positioned on each side of the monitor. Visual stimuli were presented on an LCD screen with a 60 Hz refresh rate (EIZO FlexScan S2202W). Button presses were recorded using a standard keyboard. Participants were seated in front of the monitor and loudspeakers at a distance of 85 cm in an electrically shielded, sound-attenuated room.

*Bayesian modelling analysis for behavioral data*

The Bayesian modelling analysis fitted several competing computational models to the individual behavioral numeric reports. Briefly, the generative model of the BCI model (Fig. 3A) assumes that common (*C*=1) or independent (*C*=2) causes are determined by sampling from a binomial distribution with the causal prior *p*(*C*=1) = *p*_common_ (i.e., prior “binding tendency”) (Odegaard & Shams, 2016). For a common cause, the “true” number of audiovisual stimuli *N*_AV_ is drawn from the numeric prior distribution N(*μ*_P_, *σ*_P_). For two independent causes, the “true” auditory (*N*_A_) and visual (*N*_V_) numbers of stimuli are drawn independently from this numeric prior distribution. Sensory noise is introduced by drawing the sensory percepts *x*_A_ and *x*_V_ independently from normal distributions centered on the true auditory (respectively visual) number of stimuli with parameters *σ*_A_ (respectively *σ*_V_). Thus, the basic generative model included the following free parameters: the causal prior *p*_common_, the numeric prior’s mean *μ*_P_ and standard deviation *σ*_P_, the auditory standard deviation *σ*_A_, and the visual standard deviation *σ*_V_. Given the sensory percepts *x*_A_ and *x*_V_ , the observer infers the posterior probability of the underlying causal structure by combining the causal prior with the sensory evidence according to Bayes rule:

$$\left( 2 \right) p\left( C=1 | x\text{A},x\text{V} \right)=\frac{p(x\text{A},x\text{V}\text{|C=1})p\text{common}}{p(x\text{A},x\text{V})}$$

The causal prior quantifies observers’ prior belief that flashes and beeps arise from a common cause and should hence be integrated. In the case of a common cause (*C*=1), the optimal audiovisual numeric estimate ($\hat{N}\text{AV,C=1}$) is obtained by combining the auditory and visual numeric percepts as well as the numeric prior weighted by their relative precisions:

$$\left( 3 \right) \hat{N}\text{AV,C=1}=\frac{\frac{x\text{A}}{\sigma\text{A}\text{2}}+\frac{x\text{V}}{\sigma\text{V}\text{2}}+\frac{\mu\text{P}}{\sigma\text{P}\text{2}}}{\frac{1}{\sigma\text{A}\text{2}}+\frac{1}{\sigma\text{V}\text{2}}+\frac{1}{\sigma\text{P}\text{2}}}$$

In the case of independent causes (*C*=2), the optimal numeric estimates of the unisensory auditory ($\hat{N}\text{A,C=2}$) and visual ($\hat{N}\text{V,C=2}$) stimuli are independent:

$$\left( 4 \right) \hat{N}\text{A,C=2}=\frac{\frac{x\text{A}}{\sigma\text{A}\text{2}}+\frac{\mu\text{P}}{\sigma\text{P}\text{2}}}{\frac{1}{\sigma\text{A}\text{2}}+\frac{1}{\sigma\text{P}\text{2}}}, \hat{N}\text{V,C=2}=\frac{\frac{x\text{V}}{\sigma\text{V}\text{2}}+\frac{\mu\text{P}}{\sigma\text{P}\text{2}}}{\frac{1}{\sigma\text{V}\text{2}}+\frac{1}{\sigma\text{P}\text{2}}}$$

To provide a final estimate of the number of signals that takes the causal uncertainty into account, the BCI model formally requires that the three numeric estimates ($\hat{N}\text{AV,C=1}$, $\hat{N}\text{A,C=2}$, $\hat{N}\text{V, C=2}$) are combined into the final task-relevant numeric estimate ($\hat{N}\text{A}$ or $\hat{N}\text{V}$) depending on the posterior probabilities of the estimates’ underlying causal structures. The observer may combine the numerical estimates according to different decision strategies (Wozny et al., 2010):

In the ‘model averaging’ strategy, the observers weighs the estimates in proportion to the posterior probabilities of their underlying causal structures:

$$\left( 5 \right) \hat{N}\text{A }\text{= p(C=1|}x\text{A},x\text{V}\text{)} \hat{N}\text{AV,C=1 }\text{+ (1 - p(C=1|}x\text{A},x\text{V}\text{)} )\hat{N}\text{A,C=2 }\text{ }$$

$$\hat{N}\text{V }\text{= p(C=1|}x\text{A},x\text{V}\text{)} \hat{N}\text{AV,C=1 }\text{+ (1 - p(C=1|}x\text{A},x\text{V}\text{)} )\hat{N}\text{V,C=2 }\text{ }$$

In the model selection strategy, the observer selects the numeric estimate that has a higher posterior probability:

$$\left( 6 \right) \hat{N}\text{A }\text{= }\left\{ \begin{aligned} \hat{N}\text{A,C=1}\text{ }\text{if}\text{ }\text{p(C=1|}x\text{A},x\text{V}\text{)}> 0.5 \\ \hat{N}\text{A,C=2}\text{ }\text{if}\text{ }\text{p(C=1|}x\text{A},x\text{V}\text{)}\leq0.5 \end{aligned} \right. , \hat{N}\text{V}\text{ }\text{= }\left\{ \begin{aligned} \hat{N}\text{V,C=1}\text{ }\text{if}\text{ }\text{p(C=1|}x\text{A},x\text{V}\text{)}> 0.5 \\ \hat{N}\text{V,C=2}\text{ }\text{if}\text{ }\text{p(C=1|}x\text{A},x\text{V}\text{)}\leq0.5 \end{aligned} \right.$$

In probability matching, the observer (non-optimally) selects the numerical estimate stochastically in proportion to the posterior causal probabilities:

$$\left( 7 \right) \hat{N}\text{A }\text{= }\left\{ \begin{aligned} \hat{N}\text{AV,C=1}\text{ if}\text{ }\text{p(C=1|}x\text{A},x\text{V}\text{)}> \alpha\\ \hat{N}\text{A,C=2}\text{ if}\text{ }\text{p(C=1|}x\text{A},x\text{V}\text{)}\leq\alpha\end{aligned} \right., \hat{N}\text{V}\text{ }\text{= }\text{ }\left\{ \begin{aligned} \hat{N}\text{AV,C=1}\text{ if}\text{ }\text{p(C=1|}x\text{A},x\text{V}\text{)}> \alpha\\ \hat{N}\text{V,C=2}\text{ if}\text{ }\text{p(C=1|}x\text{A},x\text{V}\text{)}\leq\alpha\end{aligned}, \alpha\sim U\left( 0,1 \right) \right.$$

We compared these three decisional strategies with two additional heuristic models (Acerbi et al., 2018). In a stochastic-fusion model, observers stochastically choose the audiovisual numeric (i.e. fusion) or the task-relevant unisensory (i.e. segregation) estimate with a fixed probability parameter η:

$$\left( 8 \right) \hat{N}\text{A }\text{= }\left\{ \begin{aligned} \hat{N}\text{AV,C=1}\text{ if }\eta>\alpha\\ \hat{N}\text{A,C=2}\text{ if}\text{ }\eta\leq\alpha\end{aligned} \right., \hat{N}\text{V}\text{ }\text{= }\text{ }\left\{ \begin{aligned} \hat{N}\text{AV,C=1}\text{ if}\text{ }\eta>\alpha\\ \hat{N}\text{V,C=2}\text{ if}\text{ }\eta\leq\alpha\end{aligned}, \alpha\sim U\left( 0,1 \right) \right.$$

The stochastic-fusion model encompasses a potential ‘forced’ fusion (η = 1) or ‘forced’ segregation (η = 0) as well as any intermediate response behavior (0 < η < 1). Importantly, the stochastic-fusion model responds independently from the audiovisual signals’ disparity (i.e., a non-causal model) while the BCI model takes numeric disparity into account to perform the causal inference (cf. Fig. 3A). Finally, the fixed-criterion model reports the fusion- or segregation-estimate whenever the signals’ absolute numeric disparity is below or above a fixed criterion k. Thus, the fixed-criterion model implements a response heuristic without formally computing the posterior probability of the causal structure as the BCI model:

$$\left( 9 \right) \hat{N}\text{A }\text{= }\left\{ \begin{aligned} \hat{N}\text{AV,C=1}\text{ if }\left| x\text{A}-x\text{V} \right|<k \\ \hat{N}\text{A,C=2}\text{ if }\left| x\text{A}-x\text{V} \right|\geq k \end{aligned} \right., \hat{N}\text{V}\text{ }\text{= }\text{ }\left\{ \begin{aligned} \hat{N}\text{AV,C=1}\text{ if }\left| x\text{A}-x\text{V} \right|<k \\ \hat{N}\text{V,C=2}\text{ if }\left| x\text{A}-x\text{V} \right|\geq k \end{aligned} \right.$$

For the second model factor, we implemented two sensory noise models: Sensory variances were either constant across stimulus numbers or increased with stimulus number proportional to an incremental sensory variance parameter Δσ:

$$\left( 10) \sigma\text{A}^{'}=\sigma\text{A}\text{ }\text{+ △}\sigma\text{A}\text{ (S}\text{A}\text{ - 1} \right), \sigma\text{V}^{'}=\sigma\text{V}\text{ }\text{+ △}\sigma\text{V}\text{ (S}\text{V}\text{ - 1)}$$

This additional increase in variance with stimulus number incorporates the notion of scalar variability. Additionally, all models included a lapse rate parameter L to account for random responses independent of audiovisual inputs.

We fitted each of the ten models in our 5 decisional-strategies x 2 sensory-noise factorial model space to the numeric reports individually for each participant (see Rohe et al., 2019). For more efficient parameter estimation, we fitted models jointly to unisensory and audiovisual conditions. Thus, the auditory/visual noise, prior spatial variance and the lapse rate were jointly informed by unisensory and audiovisual conditions.

To obtain maximum likelihood estimates for the eight parameters of the models (*p*_common_ / k / η, *µ*_P_, *σ*_P_, *σ*_A_, *σ*_V_, $\text{△}\sigma\text{A}$, $\text{△}\sigma\text{V}$, L), we used a Bayesian optimization algorithm as implement in the BADS toolbox (Acerbi & Ma, 2017). This optimization algorithm was initialized with 50 different random parameters. We report the results (i.e., model comparisons and parameters) for models with the highest log likelihood across these initializations. Model recovery showed that the parameters could be reliably identified with very small biases using this fitting procedure (Supporting Fig. S5).

To identify the optimal model for explaining participants’ data, we compared the 10 candidate models using the Bayesian Information Criterion (BIC) as an approximation to the model evidence (Raftery, 1995). We performed Bayesian model comparison (Rigoux et al., 2014) at the random-effects group level, separately for HC and SCZ, as implemented in SPM12 (Friston et al., 1994) to obtain the protected exceedance probability (the probability that a given model is more likely than any other model, beyond differences due to chance (Rigoux et al., 2014)) for each of the 10 candidate models. Further, we performed factorial Bayesian model selection on the two-factorial model space using robust Bayesian model-selection procedures (L. Acerbi, personal communication) to compute protected exceedance probabilities for each family across the two factors in our 2 x 5 factorial model space (Fig. 3B). To assess whether the frequencies of the optimal model differ between HC and SCZ, we applied between-group Bayesian model comparison (Rigoux et al., 2014) as implemented in the variational Bayesian approach toolbox (Daunizeau et al., 2014). Using the posterior probability that the two groups have the same model frequencies, we computed the Bayes factor BF_10_ quantifying the evidence in favor of different (H1) or equivalent (H0) model frequencies (i.e., assuming equal prior probabilities).

To investigate whether HC and SCZ differ in parameters of the winning model (i.e., the BCI model with ‘model averaging’ and increasing sensory variances), we compared the parameters between groups using a two-sample two-sided randomization test (n = 5000 randomizations; Tab. 2). Further, we correlated the parameters with SCZ patients’ positive and negative symptoms as well as general psychopathy as measured with the PANSS using a randomization test of the correlation (n = 5000 randomizations).

To investigate whether HC and SCZ differentially adjusted the causal priors depending on the stimulus history, we sorted the current trials according to the previous trial’s audiovisual numeric disparity (i.e., |n_A_-n_V_| ∈ {0-3}) and selectively refitted the causal prior of the BCI model. We entered the causal prior *p*_common_ into a 4 (numeric disparity on previous trial: 0, 1, 2, 3) x 2 group (HC vs. SCZ) mixed-model ANOVA. Likewise, we investigated how observers adapted the numeric prior by sorting the current trials according to previous trials task-relevant auditory/visual signals (i.e., n_A_ or n_V_ ∈ {1-4}) and selective refitting of the numeric prior (i.e. numeric prior’s mean *μ*_P_ and standard deviation *σ*_P_; Fig. 5). We entered the numeric prior’s mean *μ*_P_ and standard deviation *σ*_P_ into separate 4 (number of task-relevant signals on previous trial: 1, 2, 3, 4) x 2 group (HC vs. SCZ) mixed-model ANOVAs. To provide evidence for the null-hypothesis i.e. no difference between groups, these model parameters were also entered into separate Bayesian mixed-model ANOVAs (see statistical analyses below).

To generate predictions for the behavioral CMB index and EEG analyses (see below) based on the BCI model, we simulated new *x*_A_ and *x*_V_ for 10000 trials for each of the 32 conditions using the fitted BCI model parameters of each participant (i.e., BCI model with model averaging and increasing sensory variances). For each simulated trial, we computed the BCI model’s i. unisensory visual ($\hat{N}\text{V,C=2}$), ii. unisensory auditory ($\hat{N}\text{A,C=2}$) estimates, iii. forced-fusion ($\hat{N}\text{A}\text{V,C=1}$), iv. final BCI audiovisual numeric estimate ($\hat{N}\text{A}$ or $\hat{N}\text{V}$ depending on whether the auditory or visual modality was task-relevant). Next, we used the mode of the resulting (kernel-density estimated) distributions for each condition and participant to compute the model predictions for the CMB index (Fig. 1D) and decoding the BCI model’s estimates from EEG patterns (see multivariate EEG analysis; Fig. 6).

*Statistical analyses of behavioral data*

We used randomization tests to analyse behavioural measures and parameters (i.e., neuropsychological test scores, CMB and BCI model parameters; Tab. S1 and Tab. 2) between conditions and groups as well as correlations between BCI model parameters and symptom scores. Randomization tests were complemented by Bayes factors BF_10_ quantifying evidence in favor of a condition or group difference or correlation (H1) relative to the null hypothesis of no difference or correlation (H0) using the bayesFactor toolbox (Krekelberg, 2021). Bayesian t-tests and correlations assumed a Jeffrey-Zellner-Siow prior and t-tests assumed a scaling factor s = 0.707 of the Cauchy prior on effects (Wetzels & Wagenmakers, 2012).

To evaluate whether experimental factors influenced HC and SCZ differentially or equivalently (e.g., for CMB), we complemented classical with Bayesian mixed-model ANOVAs (Tab. 1 and 3) with multivariate Cauchy priors on the effects and uniform model priors. Bayesian ANOVAs allow to compute inclusion Bayes factors (BF_incl_) for each experimental factor which quantify the average evidence that including the factor in the ANOVA models improves the models’ fit given a higher complexity of these models (van den Bergh et al., 2020; Wagenmakers, Love, et al., 2018). For Bayesian analyses, the following interpretations of BF apply: BF > 3 or > 10 provide substantial or strong evidence (Kass & Raftery, 1995; Wagenmakers, Marsman, et al., 2018) for condition/group differences, correlations or inclusion of a factor, whereas BF < 1/3 or < 1/10) provides substantial or strong evidence for condition/group equivalence, no correlation or exclusion the factor (van den Bergh et al., 2020). All classical and Bayesian ANOVAs were computed using JASP 0.17.2.1 (Wagenmakers, Love, et al., 2018).

*EEG – Data acquisition and preprocessing.*

EEG signals were recorded from 64 active electrodes positioned in an extended 10–20 montage using electrode caps (actiCap, Brain Products, Gilching, Germany) and two 32 channel DC amplifiers (BrainAmp, Brain Products). Electrodes were referenced to FCz using AFz as ground during recording. Signals were digitized at 1000 Hz with a high-pass filter of 0.1 Hz. Electrode impedances were kept below 25 kOhm. Preprocessing of EEG data was performed using Brainstorm 3.462 running on Matlab R2015b. EEG data were band-pass filtered (0.25–45 Hz). Eye blinks were automatically detected using data from the FP1 electrode (i.e., a blink was detected if the band-pass (1.5–15 Hz) filtered EEG signal exceeded two times the STD; the minimum duration between two consecutive blinks was 800 ms). Signal-space projectors (SSPs) were created from band-pass filtered (1.5–15 Hz) 400 ms segments centered on detected blinks. The first spatial component of the SSPs was then used to correct blink artifacts in continuous EEG data. Further, all data were visually inspected for artifacts from blinks (i.e. residual blink artifacts after correction using SSPs), saccades, motion, electrode drifts or jumps and contaminated segments were discarded from further analysis. On average, we discarded 6.171 ± 0.923 % SEM of all trials for HC and 7.216 ± 1.444 % for SCZ; the difference was not significant, t_38_ = -0.638, p = 0.528, d = -0.204, BF_10_ = 0.366. Finally, EEG data were re-referenced to the average of left and right mastoid electrodes and downsampled to 200 Hz. For analysis of event-related potentials (ERPs) and decoding analyses (see below), all EEG data were baseline corrected with a 200 ms prestimulus baseline and were analyzed from 100 ms before stimulus onset up to 750 ms after stimulus onset, when the response screen was presented.

*Analysis of response tendencies in numeric reports*

In supplemental analyses of participants’ behavioral data, we analyzed the auditory and visual numeric reports more comprehensively between groups across the combination of audiovisual signals (Supplemental Fig. S1). Both HC and SCZ underestimated higher signal numbers with increasing variances, a key prediction of the scalar-variability model of numerosity estimation (Dehaene, 2007; Gallistel & Gelman, 2000). SCZ appeared to show a stronger central tendency with stronger underestimation of large (Fig. 2A) and overestimation of small numbers (Fig. 2B). To quantify and compare the strength of participants’ central tendency, we modelled the influence of the auditory and visual signal number and their interaction on numeric reports with linear-logarithmic regressions (Supplemental Fig. S1B). Yet, slope estimates, as indicators of central tendency, did not significantly differ between both groups (Tab. S2). Significant crossmodal and interaction terms indicated that observers’ numeric reports where biased by the task-irrelevant signals.

*Decoding of group membership from EEG response patterns*

In supplemental multivariate analyses of EEG data, we assessed whether schizophrenia more generally changes the multivariate ERP response patterns (i.e., grand average ERPs across electrones) of basic sensory components and early audiovisual interactions (Supplemental Fig. S4). If multivariate response patterns differed systematically between HC and SCZ, a multivariate decoder would be able to distinguish between both groups based on the response patterns (Koch et al., 2015). Thus, we trained a linear support-vector machine classification (SVC) using LibSVM (Chang & Lin, 2011) to classify the diagnostic group from ERP patterns in all but one participant. The trained SVC then predicted the group from EEG patterns of the left-out participant. In a leave-one-participant-out crossvalidation scheme (Koch et al., 2015), the training-test procedure was repeated for all participants and decoding accuracy was computed as the fraction of correct classifications across all participants. The SVC’s parameter ν was optimized using a grid search within each cross-validation fold (i.e., nested cross-validation). This training-test procedure was repeated for all 20 ms time windows (i.e., 64 channels x 4 time points = 256 features) and unisensory as well as audiovisual ERP patterns. To test whether response patterns represented information on group membership (i.e., the SVC’s decoding accuracies exceeded chance level), we used a non-parametric randomization test (5000 randomizations) in which we computed the decoding accuracy of the SVC-predicted group membership as test statistic. In each randomization fold, we applied the SVC’s crossvalidation scheme on randomized group membership. To correct for multiple comparisons across the EEG sampling points, we used a cluster-based correction (Maris & Oostenveld, 2007) with the sum of the decoding accuracies across a cluster as cluster-level statistic and an auxiliary cluster-defining threshold of decoding accuracy = 0.55 for each time point. However, the decoder was not able to predict group membership from unisensory or audiovisual congruent ERP patterns (p > 0.05) as decoding accuracies did not exceed chance level for any time points because ERP activation patterns were highly correlated across both groups (Supplemental Fig. S4).

**Supporting Information Figures and Tables**

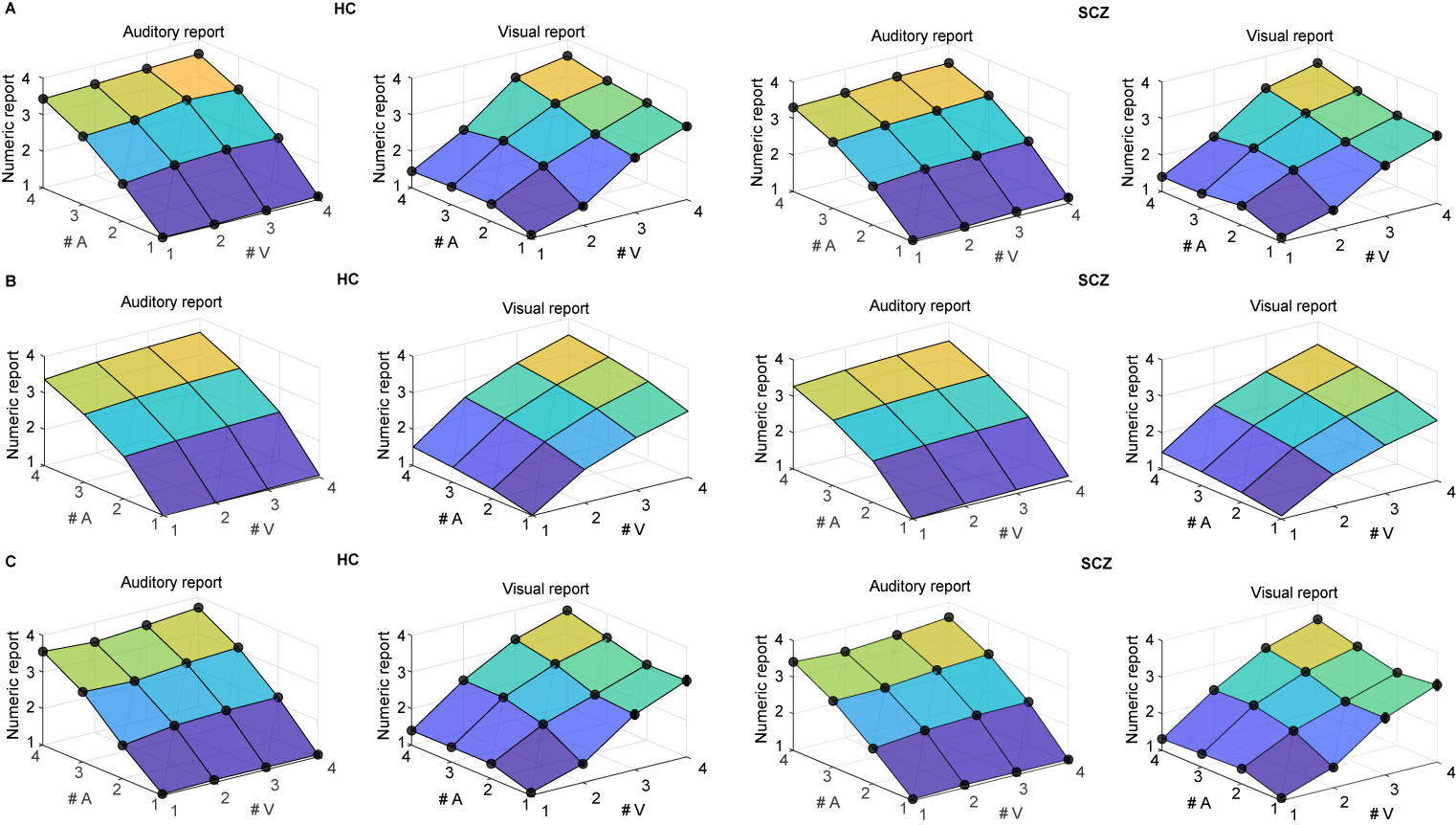

**Figure S1. Numeric reports and predictions from a log-linear regression model as well as the BCI model as a function of the visual signal number, auditory signal number and task, separately for HC (left two columns, n = 23) and SCZ (right two columns, n = 17)**. **(A)** Mean numeric reports (across participants mean ± SEM). **(B)** Mean predictions (across-participants mean) from a log-linear regression model that predicted the numeric reports from the logarithmic visual and auditory signal numbers as well as their interaction (i.e., r_A/V_ = b_A_ * log(n_A_) + b_V_ * log(n_V_) + b_AxV_ * log(n_V_) * log(n_A_) + c). **(C)** Mean predictions from the BCI model (across-participants mean ± SEM; model-averaging with increasing sensory variances) reproduced the logarithmic compression of numeric reports.

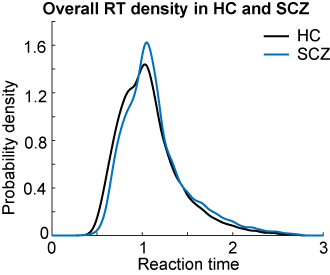

**Figure S2. Overall reaction time (RT) density in HC and SCZ (n = 40).** When fitting ex-Gaussian functions to the RT distributions, the functions’ mean (t_38_ = -1.325, p = 0.193, d = -0.424, BF_10_ = 0.620), variance (t_38_ = -0.511, p = 0.612, d = -0.164, BF10 = 0.346) and lambda parameter (t_38_ = -0.607, p = 0.548, d = -0.194, BF10 = 0.361) were not significantly different, but rather equivalent, between both groups.

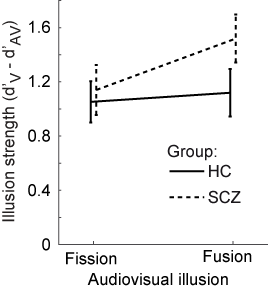

**Figure S3. The strength of the fission and fusion illusions (across-participants mean ± SEM) is shown as a function of group (HC vs. SCZ, n = 40).** The illusions were computed as difference in sensitivity (d prime, d’) between the unisensory baseline condition (V1A0 vs. V2A0) and the illusion conditions (fission: V1A2 vs. V2A2; fusion: V1A1 vs. V2A2). For the fission or fusion conditions, response ‘2’ or ‘1’, respectively, were defined as signal. Thus, this illusion measure accounts for a possible shift in the response criterion which could be confounded with the fission or fusion illusions (Vanes et al, 2016). In contrast to Vanes et al. (2016), a mixed-model ANOVAs did not reveal a significant difference in illusion strength between the two types of illusions (factor audiovisual illusion, F_1,38_ = 2.598 p = 0.115, part. η^2^ = 0.064, BF_incl_ = 0.461) or a difference between HC and SCZ participants (factor group: F_1,38_ = 1.431, p = 0.239, part. η^2^ = 0.036, BF_incl_ = 0.440; interaction illusion × group: F_1,38_ = 1.271, p = 0.267, part. η^2^ = 0.032, BF_incl_ = 0.273). Note that we only included trials in which participants counted the number of flashes as ≤ 2.

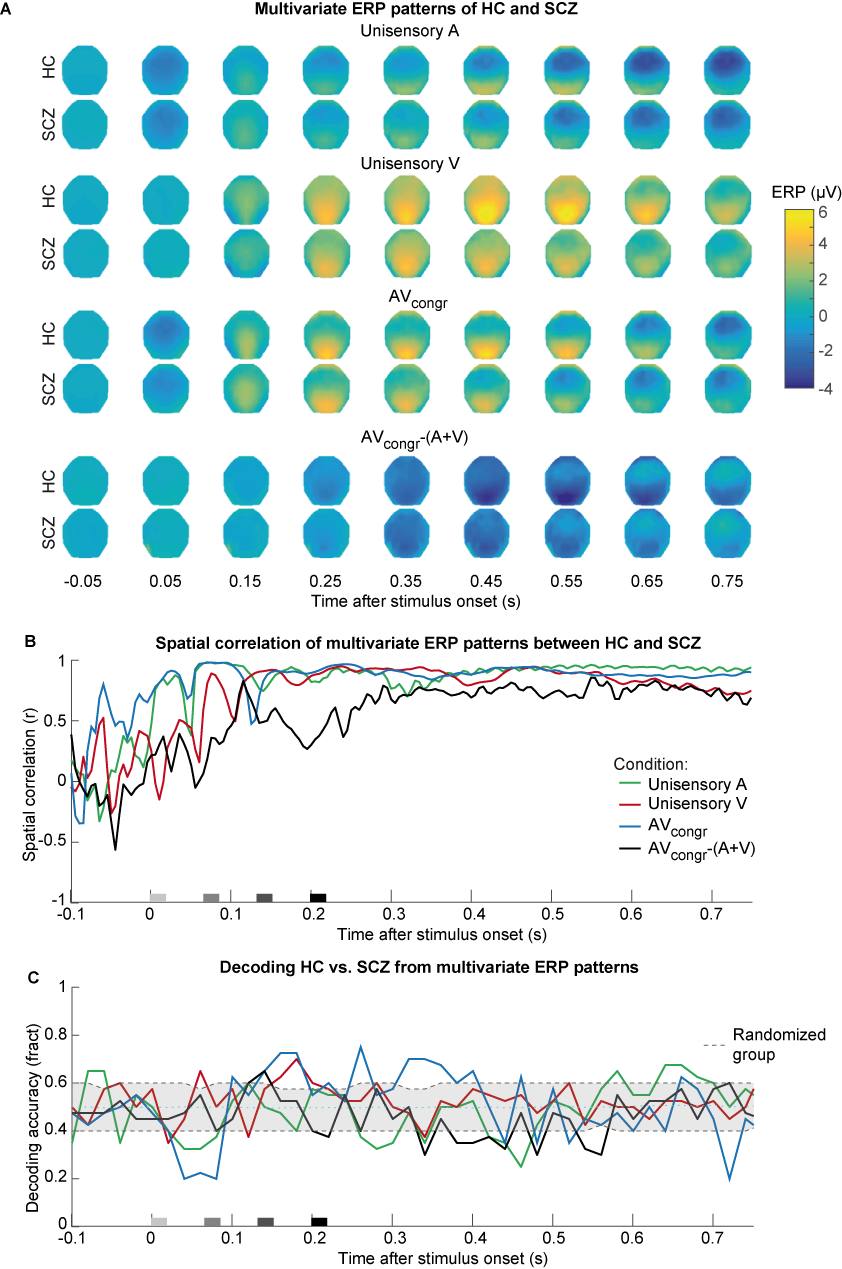

**Figure S4. Multivariate ERP patterns (i.e., topographies, the spatial correlation of patterns between HC and SCZ and decoding results (n = 40) .** **(A)** Patterns are shown as a function of time, averaged in 100 ms time windows and separately for HC and SCZ participants. **(B)** Spatial correlation of multivariate ERP patterns between HC and SCZ participants as a function of time, separately for the four conditions (N.B.: Spatial correlations before stimulus onset might arise from expectation processes). **(C)** Decoding accuracy (i.e., fraction of correct classifications) of a decoder trained to classify the participant group (i.e. HC vs. SCZ) from ERP patterns of the four conditions. No significant clusters (p > 0.05) of decoding accuracies above the chance level of 0.5 were found in one-sided cluster-based corrected randomization tests. As a reference, the grey dashed lines indicate decoding accuracy (fraction ± 68% CI; averaged across the four conditions) of the decoder trained on randomized group membership (n = 5000 randomizations).

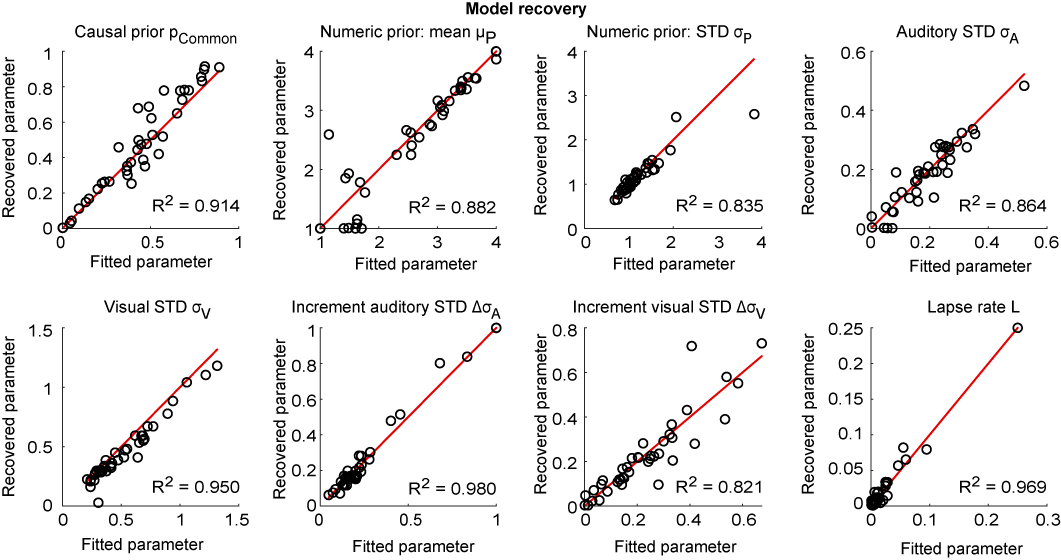

**Figure S5. Results of the model recovery (n = 40).** For model recovery, the winning model (i.e., model averaging with increasing sensory variances) predicted responses which were then again fitted to obtain recovered parameters. The plots show the recovered parameters as a function of the parameters originally fitted to participants’ behavioral data. The red line is a line with slope 1 and intercept 0.

|  | **Table S1. Demographic, psychopathological and neuropsychological data (across-participants mean ± STD) from healthy controls (HC) and participants with schizophrenia (SCZ).** | | | | | |
| --- | --- | --- | --- | --- | --- | --- |
|  | | **HC** | **SCZ** | **t_38_** | **p** | **BF_10_** |
| n | | 23 | 17 | - | - | - |
| Age (years) | | 35.96±2.50 | 33.12±1.99 | 0.84 | 0.414 | 0.412 |
| Sex (% w) | | 43.5 | 17.6 | 2.97(χ2) | 0.085 | - |
| Education (years) | | 16.65±0.63 | 15.44±0.89 | 1.14 | 0.269 | 0.520 |
| PANSS Pos. Sympt. (Σ items) | | - | 13.59±0.97 | - | - | - |
| PANSS Neg. Sympt. (Σ items) | | - | 17.29±1.75 | - | - | - |
| PANSSGen | | - | 28.88±1.71 | - | - | - |
| CPZEquiv | | - | 572.59±62.95 | - | - | - |
| VLMT free recall (Σ trial 1-5) | | 57.83±1.65 | 49.41±2.99 | 2.63 | 0.009 | 4.230 |
| VLMT delayed recall (trial 7) | | 12.61±0.56 | 10.47±0.64 | 2.49 | 0.018 | 3.287 |
| VLMT recognition (Hits-FA) | | 13.87±0.32 | 12.91±0.54 | 1.61 | 0.104 | 0.853 |
| TMT A (sec) | | 28.60±1.98 | 25.70±2.99 | 0.84 | 0.406 | 0.413 |
| TMT B (sec) | | 59.89±3.09 | 67.75±10.72 | -0.80 | 0.462 | 0.401 |
| MWT-B (raw value) | | 30.78±0.67 | 27.57±1.32 | 2.34 | 0.019 | 2.523 |
| Stroop effect (sec) | | 28.93±2.34 | 33.61±3.49 | -1.16 | 0.258 | 0.527 |
| EHI (Laterality index) | | 58.19±10.65 | 69.42±8.23 | -0.79 | 0.439 | 0.398 |
| BDI (Σ items) | | 2.09±0.62 | 13.47±1.68 | -7.05 | <0.001 | >100 |
| CDSS (⌀ items) | | - | 0.33±0.08 | - | - | - |
| Note: PANSS: Positive and Negative Symptom Scale (with three subscales: Positive Symptoms/Pos. Sympt., Negative Symptoms/Neg. Sympt. and General Psychopathology/PANSSGen); CPZEquiv: Chlorpromazine equivalents (mg); VLMT: Verbal learning and memory test; TMT: Trail Making Test; MWT-B: Mehrfachwahl-Wortschatz-Intelligenztest (Multipe word choice intelligent test); Stroop effect: Time color word naming – time color plate naming; EHI: Edinburgh Handedness Inventory; BDI: Beck’s Depression Inventory; CDSS: Calgary Depression Scale for Schizophrenia with 9 subscales | | | | | | |

| **Table S2. Statistical significance of regression parameter estimates across and between HC and SCZ in the log-linear regression model that predicted the numeric reports from the visual and auditory signal numbers as well as their interaction.** | | | | | |
| --- | --- | --- | --- | --- | --- |
| **One-sample t test on parameter estimates across HC and SCZ** | | | | | |
| Task relevance | Parameter estimates | t | p | d | BF_10_ |
| A report | b_A_ | 48.770 | <0.001 | 7.711 | >100 |
|  | b_V_ | 4.941 | <0.001 | 0.781 | >100 |
|  | b_AxV_ | 3.265 | 0.002 | 0.516 | 14.687 |
| V report | b_A_ | 6.055 | <0.001 | 0.957 | >100 |
|  | b_V_ | 18.777 | <0.001 | 2.969 | >100 |
|  | b_AxV_ | 3.406 | 0.002 | 0.539 | 20.812 |
| **Mixed-effects ANOVA on parameter estimates** | | | | | |
| Effect | F | df1, df2 | p | part. η^2^ | BF_Incl_ |
| TR | 4.853 | 1, 38 | 0.034 | 0.113 | >100 |
| PE | 185.289 | 3, 38 | <0.001 | 0.830 | >100 |
| Group | 2.250 | 1, 38 | 0.142 | 0.056 | 0.115 |
| TRxPE | 288.072 | 2.149, 81.667 | <0.001 | 0.883 | >100 |
| TRxGroup | 0.484 | 1, 38 | 0.491 | 0.013 | 0.095 |
| PExGroup | 1.448 | 3, 114 | 0.233 | 0.037 | 0.120 |
| TRxPExGroup | 0.975 | 2.149, 81.667 | 0.387 | 0.025 | 0.054 |
| Note: Regression model: r_A/V_ = b_A_ * log(n_A_) + b_V_ * log(n_V_) + b_AxV_ * log(n_V_) * log(n_A_) + c, r_A/V_ = numeric auditory/visual report, n_A_ = auditory signal number, n_V_ = visual signal number, c = constant. Parameter estimates were tested against zero across HC and SCZ using one-sample t-tests. Parameter estimates were compared between groups using a mixed-model and Bayesian ANOVA with factors TR = task relevance (auditory vs. visual report; within-participant), PE = parameter estimate from regression model (b_A_, b_V_, b_AxV_, c; within-participant) and Group (HC vs. SCZ; between-participants). Degrees of freedom are Greenhouse-Geisser corrected if sphericity is violated for an effect. | | | | | |

| **Table S3. Results of the Bayesian model comparison of 2 x 5 factorial model space with factor ‘decision strategy’ and ‘sensory variance’ in HC and SCZ participants.** | | | | | | | | | | | | | | | | |
| --- | --- | --- | --- | --- | --- | --- | --- | --- | --- | --- | --- | --- | --- | --- | --- | --- |
| Decision strategy | ***σ*** |  | ***p*_common_** | ***µ*_P_** | ***σ*_P_** | ***σ*_A_** | ***σ*_V_** | Δ***σ*_A_** | Δ***σ*_V_** | ***L*** | ***k*** | η | ***R*^2^** | **relBIC** | **pEP** | **% win** |
| MA | c | HC | 0.47±0.05 | 2.06±0.23 | 1.80±0.23 | 0.49±0.02 | 0.79±0.06 | - | - | 0.01±0.00 | - | - | 0.87±0.01 | 385.463 | 0.00 | 0 |
|  |  | SCZ | 0.49±0.07 | 1.76±0.20 | 1.18±0.09 | 0.52±0.07 | 0.81±0.13 | - | - | 0.02±0.01 | - | - | 0.87±0.03 | 271.046 | 0.00 | 0 |
| MS | c | HC | 0.46±0.05 | 1.94±0.19 | 1.66±0.23 | 0.50±0.03 | 0.76±0.05 | - | - | 0.01±0.00 | - | - | 0.87±0.01 | 104.642 | 0.00 | 0.04 |
|  |  | SCZ | 0.47±0.07 | 1.78±0.20 | 1.08±0.09 | 0.52±0.07 | 0.79±0.14 | - | - | 0.02±0.01 | - | - | 0.86±0.03 | 103.438 | 0.00 | 0 |
| PM | c | HC | 0.44±0.05 | 2.04±0.20 | 1.59±0.22 | 0.48±0.02 | 0.73±0.05 | - | - | 0.01±0.00 | - | - | 0.87±0.01 | 212.520 | 0.00 | 0 |
|  |  | SCZ | 0.46±0.07 | 1.82±0.19 | 1.09±0.09 | 0.50±0.07 | 0.76±0.12 | - | - | 0.02±0.01 | - | - | 0.87±0.03 | 184.136 | 0.00 | 0 |
| FC | c | HC | - | 2.03±0.19 | 1.50±0.21 | 0.49±0.03 | 0.73±0.05 | - | - | 0.01±0.01 | 1.34±0.13 | - | 0.87±0.01 | 131.296 | 0.00 | 0 |
|  |  | SCZ | - | 1.80±0.19 | 1.09±0.09 | 0.51±0.06 | 0.82±0.16 | - | - | 0.02±0.01 | 1.16±0.18 | - | 0.87±0.03 | 135.270 | 0.00 | 0 |
| SF | c | HC | - | 2.05±0.22 | 1.91±0.26 | 0.47±0.03 | 0.71±0.05 | - | - | 0.01±0.00 | - | 0.22±0.03 | 0.87±0.01 | 172.262 | 0.00 | 0 |
|  |  | SCZ | - | 1.74±0.21 | 1.15±0.11 | 0.47±0.07 | 0.72±0.13 | - | - | 0.02±0.01 | - | 0.25±0.04 | 0.87±0.03 | 155.330 | 0.00 | 0 |
| MA | i | HC | 0.41±0.05 | 2.61±0.22 | 1.36±0.13 | 0.20±0.02 | 0.54±0.06 | 0.25±0.04 | 0.25±0.04 | 0.01±0.00 | - | - | 0.90±0.01 | 1452.594 | 1.00 | 0.7 |
|  |  | SCZ | 0.46±0.07 | 2.47±0.22 | 1.03±0.05 | 0.17±0.03 | 0.49±0.06 | 0.27±0.07 | 0.20±0.03 | 0.03±0.02 | - | - | 0.90±0.02 | 1053.740 | 0.99 | 0.65 |
| MS | i | HC | 0.35±0.04 | 2.50±0.20 | 1.28±0.09 | 0.22±0.03 | 0.54±0.05 | 0.26±0.05 | 0.20±0.03 | 0.01±0.00 | - | - | 0.89±0.01 | 984.443 | 0.00 | 0 |
|  |  | SCZ | 0.42±0.06 | 2.30±0.20 | 1.01±0.03 | 0.20±0.03 | 0.53±0.06 | 0.24±0.07 | 0.17±0.04 | 0.03±0.01 | - | - | 0.89±0.02 | 800.861 | 0.00 | 0.18 |
| PM | i | HC | 0.30±0.05 | 2.52±0.20 | 1.26±0.08 | 0.22±0.03 | 0.54±0.06 | 0.23±0.05 | 0.21±0.04 | 0.01±0.00 | - | - | 0.89±0.01 | 997.222 | 0.00 | 0.04 |
|  |  | SCZ | 0.38±0.07 | 2.29±0.21 | 1.01±0.04 | 0.20±0.03 | 0.50±0.06 | 0.25±0.07 | 0.17±0.03 | 0.03±0.01 | - | - | 0.89±0.02 | 855.084 | 0.00 | 0 |
| FC | i | HC | - | 2.47±0.21 | 1.29±0.10 | 0.22±0.03 | 0.55±0.07 | 0.24±0.05 | 0.25±0.06 | 0.01±0.00 | 0.85±0.13 | - | 0.89±0.01 | 981.848 | 0.00 | 0.13 |
|  |  | SCZ | - | 2.31±0.21 | 1.01±0.04 | 0.20±0.03 | 0.51±0.06 | 0.25±0.07 | 0.19±0.05 | 0.03±0.01 | 0.93±0.18 | - | 0.89±0.02 | 775.682 | 0.00 | 0 |
| SF | i | HC | - | 2.60±0.23 | 1.59±0.17 | 0.20±0.03 | 0.55±0.06 | 0.24±0.05 | 0.21±0.04 | 0.01±0.00 | - | 0.19±0.03 | 0.90±0.01 | 1048.903 | 0.00 | 0.09 |
|  |  | SCZ | - | 2.35±0.22 | 1.03±0.05 | 0.17±0.03 | 0.47±0.06 | 0.25±0.07 | 0.17±0.04 | 0.03±0.01 | - | 0.21±0.04 | 0.89±0.02 | 818.691 | 0.01 | 0.18 |
| Note: *Factor decision strategies*: MA, model averaging; MS, model selection; PM, probability matching; FC, fixed-criterion model; SF, stochastic fusion model; *Factor sensory variance*: c, constant sensory variance across signal numbers; i, increasing sensory variance with larger signal numbers; *Factor model parameters*: *p*_common_, causal prior; *µ*_P_, mean of the numeric prior; *σ*_P_, standard deviation of the numeric prior; *σ*_A_, standard deviation of the auditory likelihood; *σ*_V_, standard deviation of the visual likelihood; Δ***σ*** increment of standard deviation per signal number; L, lapse parameter; k, fixed criterion threshold; η, probability of stochastic fusion; *Model fit and comparison statistics*: R^2^, Nagelkerke’s coefficient of determination (Nagelkerke, 1991) using a null model of random guesses of stimulus number 1-4 with equal probability 0.25; relBIC, Bayesian information criterion at the group level, i.e. subject-specific BICs summed over all subjects (BIC = LL − 0.5 *m* ln(n), LL = log likelihood, *m* = number of parameters, *n* = number of data points) of a model relative to the worst model (n.b. a larger relBIC indicates that a model provides a better explanation of our data); pEP, protected exceedance probability, i.e. the probability that a given model is more likely than any other model, beyond differences due to chance). % win, percentage of participants in which a model won the within-participant model comparison based on BIC. | | | | | | | | | | | | | | | | |

|  | **Table S4. Results of testing the accuracies of decoding the BCI estimates from EEG patterns against zero in HC, SCZ and their group difference.** | | | | | | | | | | | |
| --- | --- | --- | --- | --- | --- | --- | --- | --- | --- | --- | --- | --- |
|  | BCI estimate | | HC | | | SCZ | | | | HC vs. SCZ | | |
|  |  |  | time (ms) | | p | time (ms) | | p | | time (ms) | | p |
| $\hat{N}\text{V,C=2}$ | | 60-740 | | < 0.001 | | 100-660 | <0.001 | | 120-220 | | 0.029 | |
| $\hat{N}\text{A,C=2}$ | | 80-740 | | <0.001 | | 100-600  640-700 | <0.001  0.047 | |  | | n.s. | |
| $\hat{N}\text{AV,C=1}$ | | 40-740 | | <0.001 | | 80-740 | <0.001 | |  | | n.s. | |
| $\hat{N}\text{A}$ or $\hat{N}\text{V}$ | | 80-740 | | <0.001 | | 80-740 | <0.001 | |  | | n.s. | |
|  | Note: The decoded BCI model’s internal estimates comprise of: i. the unisensory visual ($\hat{N}\text{V,C=2}$), ii. the unisensory auditory ($\hat{N}\text{A,C=2}$) estimates under the assumption of independent causes (*C*=2), iii. the forced-fusion estimate ($\hat{N}\text{AV,C=1}$) under the assumption of a common cause (*C*=1) and iv. the final BCI estimate ($\hat{N}\text{A}$ or $\hat{N}\text{V}$ depending on the sensory modality that is task-relevant) that averages the task-relevant unisensory and the precision-weighted estimate by the posterior probability estimate of each causal structure $\left( p(C=1 \vert x\text{A},x\text{V} \right)$). The table denotes significant clusters in one-sided (HC, SCZ) or two-sided (HC vs. SCZ) cluster-based corrected randomization t-test. n.s. denotes no significant clusters. | | | | | | | | | | | |

**Supporting Information References**

Acerbi, L., & Ma, W. J. (2017). Practical Bayesian optimization for model fitting with Bayesian adaptive direct search. *Advances in neural information processing systems (NIPS)*, 1836-1846.

Addington, D., Addington, J., & Schissel, B. (1990). A depression rating scale for schizophrenics. *Schizophrenia research*, *3*(4), 247-251.

Bäumler, G. (1984). *Farbe-Wort-Interferenztest (FWIT) nach JR Stroop*. Hogrefe Verlag GmbH, Göttingen, Germany.

Brainard, D. H. (1997). The psychophysics toolbox. *Spatial vision*, *10*(4), 433-436.

Chang, C. C., & Lin, C. J. (2011). LIBSVM: a library for support vector machines. *ACM Transactions on Intelligent Systems and Technology (TIST)*, *2*(3), 27.

Daunizeau, J., Adam, V., & Rigoux, L. (2014). VBA: a probabilistic treatment of nonlinear models for neurobiological and behavioural data. *PLoS Comput Biol*, *10*(1), e1003441.

de Gelder, B., Vroomen, J., Annen, L., Masthof, E., & Hodiamont, P. (2003). Audio-visual integration in schizophrenia. *Schizophr Res*, *59*(2-3), 211-218. <https://doi.org/S0920996401003449> [pii]

Dehaene, S. (2007). Symbols and quantities in parietal cortex: Elements of a mathematical theory of number representation and manipulation. *Sensorimotor foundations of higher cognition*, *22*, 527-574.

Friston, K. J., Holmes, A. P., Worsley, K. J., Poline, J. P., Frith, C. D., & Frackowiak, R. S. J. (1994). Statistical parametric maps in functional imaging: a general linear approach. *Hum Brain Mapp*, *2*(4), 189-210.

Gallistel, C. R., & Gelman, I. I. (2000). Non-verbal numerical cognition: from reals to integers. *Trends Cogn Sci*, *4*(2), 59-65. <https://doi.org/10.1016/s1364-6613(99)01424-2>

Hautzinger, M., Keller, F., & Kühner, C. (2006). Das Beck Depressionsinventar II. Deutsche Bearbeitung und Handbuch zum BDI II. *Frankfurt a. M: Harcourt Test Services*.

Helmstaedter, C., Lendt, M., & Lux, S. (2001). *VLMT: Verbaler Lern-und Merkfähigkeitstest*. Beltz Test, Göttingen, Germany.

Kass, R. E., & Raftery, A. E. (1995). Bayes factors. *Journal of the american statistical association*, *90*(430), 773-795.

Kay, S. R., Fiszbein, A., & Opler, L. A. (1987). The positive and negative syndrome scale (PANSS) for schizophrenia. *Schizophr Bull*, *13*(2), 261.

Koch, S. P., Hägele, C., Haynes, J.-D., Heinz, A., Schlagenhauf, F., & Sterzer, P. (2015). Diagnostic classification of schizophrenia patients on the basis of regional reward-related FMRI signal patterns. *PLoS One*, *10*(3), e0119089.

Krekelberg, B. (2021). *bayesFactor*. Retrieved January 30th from <https://klabhub.github.io/bayesFactor/>

Lehrl, S. (2005). *Mehrfachwahl-Wortschatz-Intelligenztest MWT-B.* Spitta Verlag GmbH, Balingen, Germany.

Leucht, S., Samara, M., Heres, S., Patel, M. X., Woods, S. W., & Davis, J. M. (2014). Dose equivalents for second-generation antipsychotics: the minimum effective dose method. *Schizophr Bull*, *40*(2), 314-326.

Maris, E., & Oostenveld, R. (2007). Nonparametric statistical testing of EEG-and MEG-data. *Journal of neuroscience methods*, *164*(1), 177-190.

Nagelkerke, N. J. (1991). A note on a general definition of the coefficient of determination. *Biometrika*, *78*(3), 691-692.

Odegaard, B., & Shams, L. (2016). The brain’s tendency to bind audiovisual signals is stable but not general. *Psychological science*, *27*(4), 583-591.

Oldfield, R. C. (1971). The assessment and analysis of handedness: the Edinburgh inventory. *Neuropsychologia*, *9*(1), 97-113.

Pearl, D., Yodashkin-Porat, D., Katz, N., Valevski, A., Aizenberg, D., Sigler, M., Weizman, A., & Kikinzon, L. (2009). Differences in audiovisual integration, as measured by McGurk phenomenon, among adult and adolescent patients with schizophrenia and age-matched healthy control groups. *Compr Psychiatry*, *50*(2), 186-192. <https://doi.org/10.1016/j.comppsych.2008.06.004>

S0010-440X(08)00089-8 [pii]

Raftery, A. E. (1995). Bayesian model selection in social research. *Sociological Methodology 1995, Vol 25*, *25*, 111-163. <https://doi.org/Doi> 10.2307/271063

Reitan, R. M. (1992). *Trail Making Test: Manual for administration and scoring*. Reitan Neuropsychology Laboratory.

Rigoux, L., Stephan, K. E., Friston, K. J., & Daunizeau, J. (2014). Bayesian model selection for group studies—revisited. *Neuroimage*, *84*, 971-985.

Shams, L., Kamitani, Y., & Shimojo, S. (2000). What you see is what you hear. *Nature*, *408*(6814), 788. <https://doi.org/10.1038/35048669>

Shams, L., Ma, W. J., & Beierholm, U. (2005). Sound-induced flash illusion as an optimal percept. *Neuroreport*, *16*(17), 1923-1927.

van den Bergh, D., Van Doorn, J., Marsman, M., Draws, T., Van Kesteren, E.-J., Derks, K., Dablander, F., Gronau, Q. F., Kucharský, Š., & Gupta, A. R. K. N. (2020). A tutorial on conducting and interpreting a Bayesian ANOVA in JASP. *LAnnee psychologique*, *120*(1), 73-96.

Wagenmakers, E.-J., Love, J., Marsman, M., Jamil, T., Ly, A., Verhagen, J., Selker, R., Gronau, Q. F., Dropmann, D., & Boutin, B. (2018). Bayesian inference for psychology. Part II: Example applications with JASP. *Psychonomic bulletin & review*, *25*(1), 58-76.

Wagenmakers, E.-J., Marsman, M., Jamil, T., Ly, A., Verhagen, J., Love, J., Selker, R., Gronau, Q. F., Šmíra, M., & Epskamp, S. (2018). Bayesian inference for psychology. Part I: Theoretical advantages and practical ramifications. *Psychonomic bulletin & review*, *25*(1), 35-57.

Wetzels, R., & Wagenmakers, E. J. (2012). A default Bayesian hypothesis test for correlations and partial correlations. *Psychon Bull Rev*, *19*(6), 1057-1064. <https://doi.org/10.3758/s13423-012-0295-x>

Williams, L. E., Light, G. A., Braff, D. L., & Ramachandran, V. S. (2010). Reduced multisensory integration in patients with schizophrenia on a target detection task. *Neuropsychologia*, *48*(10), 3128-3136.

Wittchen, H.-U., Wunderlich, U., Gruschwitz, S., & Zaudig, M. (1997). SKID I. Strukturiertes Klinisches Interview für DSM-IV. Achse I: Psychische Störungen. Interviewheft und Beurteilungsheft. Eine deutschsprachige, erweiterte Bearb. d. amerikanischen Originalversion des SKID I.

Wozny, D. R., Beierholm, U. R., & Shams, L. (2010). Probability matching as a computational strategy used in perception. *PLoS Comput Biol*, *6*(8). <https://doi.org/10.1371/journal.pcbi.1000871>

e1000871 [pii]
